# LGG-1/GABARAP lipidation is dispensable for autophagy and development in *C .elegans*

**DOI:** 10.1101/2021.10.05.462725

**Authors:** Romane Leboutet, Céline Largeau, Magali Prigent, Grégoire Quinet, Manuel S. Rodriguez, Marie-Hélène Cuif, Emmanuel Culetto, Christophe Lefebvre, Renaud Legouis

**Affiliations:** Université Paris-Saclay, CEA, CNRS, Institute for Integrative Biology of the Cell (I2BC), 91198, Gif-sur-Yvette, France; INSERM U1280, 91198, Gif-sur-Yvette, France; Laboratoire de Chimie de Coordination (LCC)-CNRS, UPS, 31400, Toulouse, France

**Keywords:** Atg8, LC3, GABARAP, lipidation, ubiquitin-like, di-glycine, *C. elegans*, development, CRISPR-Cas9, *S. cerevisiae*, autophagy

## Abstract

The ubiquitin-like proteins Atg8/LC3/GABARAP are required for multiple steps of autophagy such as initiation, cargo recognition and engulfment, vesicle closure and degradation. Most of LC3/GABARAP functions are considered dependent on their post-translational modifications and addressing to membranes through a conjugation to a lipid, the phosphatidylethanolamine. Contrarily to mammals, *C. elegans* possesses single homologs of LC3 and GABARAP families, named LGG-2 and LGG-1. Using site directed mutagenesis, we inhibited the conjugation of LGG-1 to the autophagosomal membrane and generated mutants that express only cytosolic forms, either the precursor or the cleaved protein. LGG-1 is an essential gene for autophagy and development in *C. elegans*, but we discovered that its functions could be fully achieved independently of its localization to the membrane. This study reveals an essential role for the cleaved form of LGG-1 in autophagy but also in an autophagy independent embryonic function. Our data question the use of the lipidated GABARAP/LC3 as the main marker of autophagic flux and highlight the high plasticity of autophagy.

## Introduction

Macroautophagy is a highly dynamic vesicular degradation system that first sequesters intracellular components in double membrane autophagosomes and then delivers them to the lysosome. Upon induction, a flat membrane is nucleated from initiation sites, generally located on the endosplamic reticulum, which further elongates around cargoes, closes, and subsequently fuses with the lysosome. The general scheme is the successive recruitment of a series of protein complexes involved in the dynamic of the process through several steps implicating the phosphorylation of lipids, the transfer of lipids from various reservoirs, the recognition of cargoes, the tethering and the fusion. Albeit the identification of numerous actors, the complete mechanism and its regulation are still poorly understood. Moreover, an increasing amount of data indicates that many of the components forming the molecular machinery for macroautophagy are also involved in autophagy-unrelated functions. One of the key players is the ubiquitin-like protein Atg8, which in yeast is required for several steps during autophagy, such as initiation, cargo recognition and engulfment, and vesicle closure (Kirisako et al., 2000; Knorr et al., 2014; Kraft et al., 2012; Nakatogawa et al., 2007; Xie et al., 2008). There are seven isoforms of Atg8 homologs in humans defining two families the MAP-LC3 (abbreviated as LC3A-a, LC3A-b, LC3B, LC3C) and the GABARAP (GABARAP, GABARAPL1, GABARAPL2) (Shpilka et al., 2011). LC3/GABARAP proteins could have both similar and very specific functions during the autophagic flux and for the recognition of specific cargoes by selective autophagy (Alemu et al., 2012; Grunwald et al., 2020; Joachim et al., 2015; Lystad et al., 2014; Pankiv et al., 2007; Weidberg et al., 2010). LC3/GABARAP proteins can bind multiple proteins through specific motifs (LIR, LC3 interacting Region) and their interactomes are only partially overlapping (Behrends et al., 2010).

The pleiotropy of Atg8/LC3/GABARAP proteins is not restricted to autophagy but extends to numerous cellular processes like intracellular receptor trafficking, tumor suppressor, ER function, LC3-associated phagocytosis, cell motility and growth (Galluzzi and Green, 2019; Schaaf et al., 2016). Such a level of complexity entangles the study of the specific functions of human LC3/GABARAP proteins (Nguyen et al., 2016). Moreover, a series of post-translational modifications, similar to the ubiquitin conjugation, is involved in the membrane targeting of Atg8/LC3/GABARAP proteins. These proteins are initially synthesized as a precursor (P), then cleaved at their C-terminus after the invariant Glycine 116 (form I), and eventually conjugated to phosphatidylethanolamine (form II) at the membrane of autophagosomes (Figure 1A) (Kabeya et al., 2000, 2004; Scherz-Shouval et al., 2003). Structural analyses have shown that LC3/ GABARAP can adopt an open or close configuration (Coyle et al., 2002). In addition, several other post-translational modifications have been reported, like phosphorylation (Cherra et al., 2010; Herhaus et al., 2020; Wilkinson et al., 2015), deacetylation (Huang et al., 2015) ubiquitination (Joachim et al., 2017) and oligomerization (Chen et al., 2007; Coyle et al., 2002), whose effects on LC3/GABARAP function and localization are largely unknown. The subcellular localization of Atg8/LC3/GABARAP proteins is multiple, either diffuse in the cytosol and nucleus, or associated to the membrane of various compartments or the cytoskeleton (Schaaf et al., 2016).

**Figure 1.**
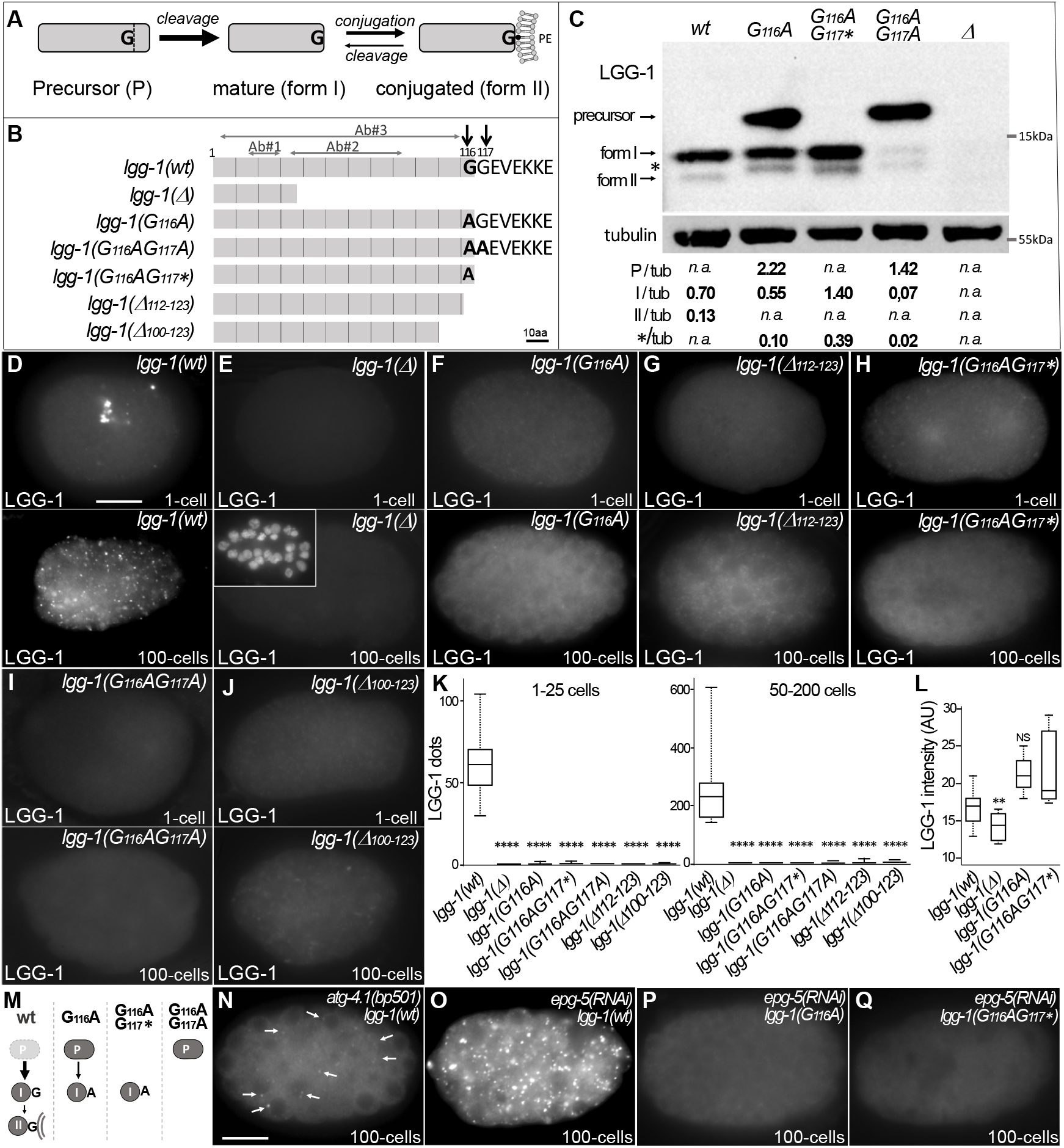
G116A abolishes the conjugation of LGG-1 to the membrane but not its cleavage. **(A)** Schematic representation of the various isoforms of Atg8s proteins after cleavage of the precursor and reversible conjugation to a phosphatidylethanolamine (PE). **(B)** Diagram of the theoretical proteins produced by the allelic *lgg-1* series used in this study. LGG-1(Δ) protein corresponds to the reference allele *lgg-1(tm3489)*, considered as a null, all others mutants have been generated using CRISP-Cas9. Black arrows point to the di-glycine residues which are mutated in alanine or stop codon (*). Other deletion mutants of the C-terminus result from non-homologous end joining. The mapping of the epitopes recognized by the LGG-1 antibodies (Ab#1, 2, 3) used in this study are indicated by horizontal grey arrows. **(C)** Western blot analysis of endogenous LGG-1 from total protein extracts of *wild-type, lgg-1(G116A), lgg-1(G116AG117*), lgg-1(G116AG117A), lgg-1(Δ)* young adults. The data shown is representative of three experiments using Ab#3 and confirmed with Ab#1. The theoretic molecular mass of the precursor, and the form I are 14.8kDa and 14.0 kDa, respectively, while the lipidated form II migrates faster. The asterisk indicates an unknown band. The quantification of each LGG-1 isoforms was normalized using tubulin. (**D-L**) Immunofluorescence analysis of endogenous LGG-1 (Ab#1 or Ab#2) in early and late embryos in *wild-type* (D), *lgg-1(Δ)* (*E*), *lgg-1(G116A)* (*F*), *lgg-1(Δ112-123)* (*G*), *lgg-1(G116AG117*)* (*H*), *lgg-1(G116AG117A)* (*I*), *lgg-1(Δ100-123)* (*J*). Inset in E show the corresponding DAPI staining of nuclei. Box-plots quantification showing the absence of puncta in all *lgg-1* mutants (K, left n= 19, 13, 11, 10, 6, 7, 6; right n= 18, 14, 12, 10, 10, 9, 12) and the increase of cytosolic staining in *lgg-1(G116A)* and *lgg-1(G116AG117*)* (L, n= 19, 13, 11, 10). Kruskal Wallis test, p-value **<0.01, ****<0.0001, NS non significant. (**M**) Schematic representation of the forms present in the wild-type and *lgg-1* mutants based on western-blot, immunofluorescence and mass spectrometry analyses. (**N-Q**) Mutant for the protease *atg-4.1(bp501)* (N) still presents few small puncta (arrows) in wild-type due to the presence of a paralog, *atg-4.2*. Autophagosome maturation defective *epg-5(RNAi)* embryos accumulate LGG-1 positive autophagosomes (O) but do not show puncta in *lgg-1(G116A)* and *lgg-1(G116AG117*)* embryos (P, Q). Scale bar is 10 μm. *(See also supplementary Figure S1 and Figure S2)*.

The existence of this versatile and pleiotropic repertoire raises a number of questions about the autophagy and the non-autophagy functions of LC3/GABARAP proteins. It is of particular interest to address the level of redundancy and specificity, including tissue specificity, of the various members, and the possible functions of the forms P and I. In the nematode *Caenorhabditis elegans*, the presence of single homologs of LC3 and GABARAP, called respectively LGG-2 and LGG-1, represents an ideal situation to characterize their multiple functions (Chen et al., 2017; Leboutet et al., 2020).

The structure of LGG-1/GABARAP and LGG-2/LC3 is highly conserved (Wu et al., 2015) and both proteins are involved in autophagy processes during development, longevity and stress (Alberti et al., 2010; Chang et al., 2017; Chen et al., 2021; Meléndez et al., 2003; Samokhvalov et al., 2008). In particular, the elimination of paternal mitochondria upon fertilization, also called allophagy (Al Rawi et al., 2011; Sato and Sato, 2011), has become a paradigm for dissecting the molecular mechanisms of selective autophagy (Djeddi et al., 2015; Zhou et al., 2016). Genetic analyses indicated that LGG-1 and LGG-2 do not have similar functions in autophagy (Alberti et al., 2010; Jenzer et al., 2019; Manil-Ségalen et al., 2014; Wu et al., 2015). During allophagy LGG-1 is important for the recognition of ubiquitinated cargoes through interaction with the specific receptor ALLO-1 (Sato et al., 2018) and the formation of autophagosomes, whereas LGG-2 is involved in their maturation into autolysosomes and trafficking (Djeddi et al., 2015; Manil-Ségalen et al., 2014). LGG-1 and LGG-2 are also differentially involved during physiological aggrephagy in embryo, with temporal-specific and cargo-specific functions (Wu et al., 2015). Based on the presence of LGG-1 and LGG-2, three populations of autophagosomes have been described in *C. elegan*s embryo: the major part are LGG-1 only, but LGG-2 only and double positives autophagosomes are also present (Manil-Ségalen et al., 2014; Wu et al., 2015). Moreover, LGG-1 is essential for embryonic and larval development, while LGG-2 is dispensable.

In this study, we investigated the functions of the non lipidated cytosolic forms of LGG-1/GABARAP. Using CRISPR-Cas9 editing, we created specific mutations in the endogenous *lgg-1* gene that result in the production of the P-like and I-like forms but not the lipidated form II. We have characterized the functionality of these new alleles of LGG-1/GABARAP for bulk autophagy, selective mitophagy and aggrephagy, but also during starvation and longevity as well as apoptotic cell engulfment and morphogenesis. LGG-1/GABARAP is an essential gene in *C. elegans*, but we discovered that most of its functions can be achieved independently of its localization to the membrane. Rescue experiments of *S. cerevisiae atg8* mutant suggest an intrinsic functionality for LGG-1 form I. We used genetic analyses to decipher whether LGG-2 could have compensatory function and show that the functionality of these LGG-1 mutants cannot be fully explained by a redundant role of LGG-2. Our data revealed that the precursor form is not capable to maintain either autophagy or developmental functions, and that its cleavage is necessary for functionality of LGG-1 form I. Further analysis of the dynamic of the autophagic flux indicated that LGG-1 form I allows the initiation and the biogenesis of autophagosomes, but form II is involved in autophagosome degradation.

## Results

### The G116G117 di-Glycine motif is a substrate for cleavage of LGG-1 precursor

The LGG-1 protein is very highly conserved from residue 1 to residue 116, sharing 92% and 74% similarity with the human GABARAP and the yeast Atg8, respectively (Manil-Ségalen et al., 2014). However, the GEVEKKE C-terminus of LGG-1 is unusual by its length and the presence of a non-conserved glycine residue in position 117 (Figure 1B, Supplementary FigS1). Found also in others Caenorhabditis species and several nematodes and arthropods, the presence of a C-terminal di-glycine is reminiscent of other ubiquitin-like proteins such as SUMO and Nedd8 (Cappadocia and Lima, 2018; Jentsch and Pyrowolakis, 2000). These specificities raise the possibility that the C-terminus could confer particular functions to the precursor and the cleaved form.

To analyze the functions of LGG-1 P and LGG-1 I, a CRISPR-Cas9 approach was used to substitute the conserved glycine 116 by an alanine, and generate three specific *lgg-1* mutants with various C-terminus (Figure 1B). In theory, both *lgg-1(G116A)* and *lgg-1(G116AG117A)* mutants are expected to accumulate a P form due to the blockage of its cleavage by ATG-4 (Wu et al., 2012). Alternatively, the *lgg-1(G116AG117*)* mutant should produce a form I. Five supplementary *lgg-1* frameshift mutants were isolated during the CRISPR experiments, resulting in deletion/insertion at the C-terminus (Figure 1B and Supplementary FigS1). Among them, *lgg-1(ΔC100-123)* and *lgg-1(ΔC112-123)* have been used later in the present study. The allele *lgg-1(tm3489)*, which deletes 48% of the open reading frame, was used as a negative control (Manil-Ségalen et al., 2014) because it is considered as a null mutant, and thereafter noted *lgg-1(Δ)*.

To assess whether *lgg-1(G116A), lgg-1(G116AG117A)* and *lgg-1(G116AG117*)* alleles code for a precursor and form I, respectively, we performed a western blot analysis with two different LGG-1 antibodies (Al Rawi et al., 2011; Springhorn and Hoppe, 2019)(Figure 1C). In basal conditions the wild-type LGG-1 is present mainly as form I (13.9 kDa) with a low amount of the faster migrating form II and no detectable precursor (14,8 kDa)(Figure 1C), while no band was observed in the allele *lgg-1(tm3489)* confirming that it is a *bona fide* null mutant.

While the *lgg-1(G116AG117*)* mutant presents a major form I, the *lgg-1(G116A)* mutant accumulates both the expected precursor form and a form I. This indicates that the cleavage of the LGG-1(G116A) precursor is still present although less efficient. In both mutant, an unexpected minor form is observed migrating differently from the lipidated form II, which is no longer detectable. The *lgg-1(G116AG117A)* mutant accumulates the precursor form (96% of the protein) indicating that the cleavage observed in the LGG-1(G116A) is dependent of the presence of a second glycine in position 117.

The substitutions in the proteins were further confirmed by mass spectrometry analyses after affinity purification of LGG-1(G116A) and LGG-1(G116AG117*) (Supplementary FigS2). The identification of C-terminal peptides validated the expected precursor form in LGG-1(G116A) and its cleavage after A116, and confirmed A116 as the last residue in LGG-1(G116AG117*). These latter forms are called hereafter “cleaved form” and “truncated form”, respectively.

### Glycine 116 is essential for lipidation of LGG-1 after cleavage

To confirm WB analyses, we next performed immunofluorescence in the embryo to analyze the subcellular localization of LGG-1 protein from the various alleles. At the 1 cell-stage and around 100 cells-stage two selective autophagy processes have been well characterized, removing paternal mitochondria and maternal aggregates, respectively (Al Rawi et al., 2011; Sato and Sato, 2011; Zhang et al., 2009). The punctate staining observed in the wild-type animals (Figure 1D) with two independent anti-LGG-1 antibodies is characteristic of the autophagosomes formed during each process, and is absent in the *lgg-1(Δ)* mutant (Figure 1E). The five mutants *lgg-1(G116A), lgg-1(G116AG117*), lgg-1(G116AG117A), lgg-1(ΔC100-123)* and *lgg-1(ΔC112-123)* present no puncta but a diffuse cytosolic staining. (Figure 1F-K), indicating that neither the precursor nor the form I are able to conjugate to the membrane The increase of the diffuse signal in *lgg-1(G116A), lgg-1(G116AG117*)* embryos (Figure 1L) suggests that the protein is less degraded in these mutants. The possibility of a small proportion of lipidated protein was further excluded by the absence of puncta after depleting the tethering factor EPG-5, which leads to the strong accumulation of autophagosomes (Figure 1O-Q)(Tian et al., 2010).

Together, these results indicate that, despite its cleavage, the G116A mutation does not allow the conjugation of LGG-1 to the membrane. LGG-1(G116AG117A) presents only a precursor form and LGG-1(G116AG117*) only a truncated form while LGG-1(G116A) produces both a precursor and a cleaved form (Figure 1M). This allelic series defines an ideal situation to address the respective roles of the precursor and form I in absence of the lipidated form II.

### Autophagy is functional in LGG-1(G116A)

To address the functionality of LGG-1 precursor and form I, we analyzed autophagy related processes that has been well characterized during *C. elegans* life cycle (Leboutet *et al*., 2020). Selective autophagy was studied in the embryonic development, during which a mitophagy and an aggrephagy process are occurring in a stereotyped manner upon fertilization and early development, respectively (Figure 2A-J). In both situations, the degradation of selective cargos was observed in the live embryo using specific labeling of the paternal mitochondria (HSP-6::GFP and mitoTracker, Figure 2A-E and Supplementary FigS3) and the P-granules (SEPA-1::GFP, Figure 2F-J). In *lgg-1(Δ)* animals, the cargos accumulate while they are degraded in the wild-type situation. In *lgg-1(G116A)*, but neither in *lgg-1(G116AG117*)* nor in *lgg-1(G116AG117A)* mutants, paternal mitochondria and SEPA-1 are degraded, suggesting that the LGG-1(G116A) protein maintains an autophagy activity (Figure 2A-K).

**Figure 2.**
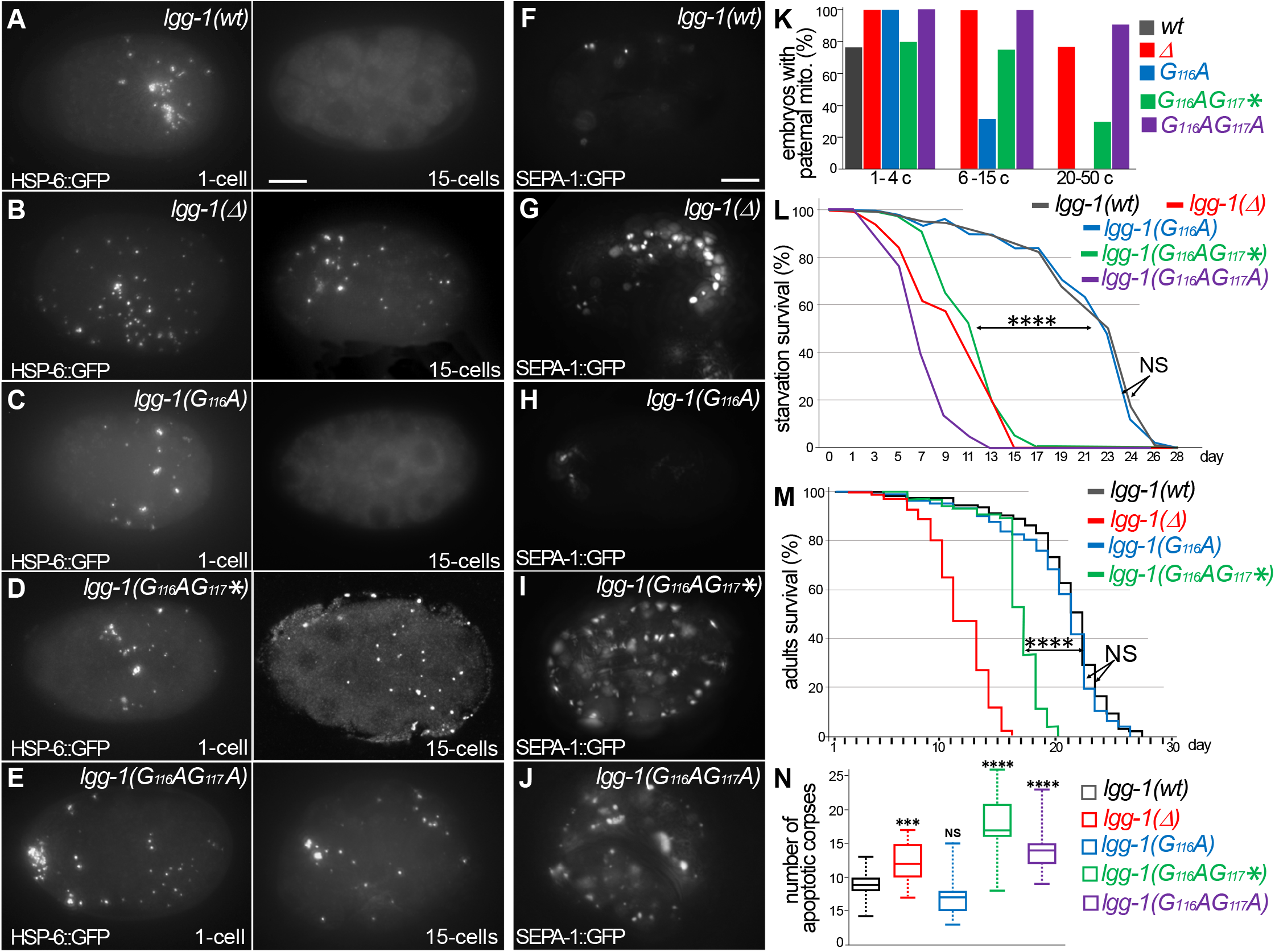
Autophagy is functional in *lgg-1(G116A)* but not in *lgg-1(G116AG117*)* and *lgg-1(G116AG117A)* (**A-E**) *In vivo* epifluorescence images of paternal mitochondria (HSP-6::GFP) at the 1-cell and 15-cells stages in *wild-type* (A), *lgg-1(Δ)* (*B*), *lgg-1(G116A)* (C), *lgg-1(G116AG117*)(D), lgg-1(G116AG117A)*(E) embryos showing an effective degradation of paternal mitochondria in *wt* and *lgg-1(G116A)* but not in *lgg-1(Δ) lgg-1(G116AG117*)* and *lgg-1(G116AG117A)*. Quantification are shown in (**K**) (**F-J**) *In vivo* epifluorescence images of the aggrephagy marker SEPA-1::GFP in late embryos for *wild-type* (F), *lgg-1(Δ)* (G), *lgg-1(G116A)* (H), *lgg-1(G116AG117*)* (I) and *lgg-1(G116AG117A)* (J). SEPA-1::GFP is degraded in *wt*, and *lgg-1(G116A)* while it accumulates in *lgg-1(Δ), lgg-1(G116AG117*)* and *lgg-1(G116AG117A)*. Scale bar is 10 μm. **(L-M)** Bulk autophagy during aging and stress was assessed by worm longevity (L, log rank test n > 100 animals, p-value ****<0.001) and starvation survival (M, Chi-square test at day 15 p-value ****<0.001). The survival is significantly reduced in *lgg-1(Δ), lgg-1(G116AG117*)* and *lgg-1(G116AG117A)* compared to *wt* and *lgg-1(G116A)*. NS non-significant. (**N**) Box-plots quantification of apoptotic corpses showing that a defective degradation in *lgg-1(G116AG117*)* and *lgg-1(G116AG117A)* but not in *lgg-1(G116A)* (n=22, 40, 46, 14, 21 Kruskal Wallis test ***<0.001, ****<0.0001, NS non-significant). *(See also supplementary Figure S3)*.

Bulk autophagy was then studied by starvation of the first stage larvae (Figure 2L). While *lgg-1(G116AG117*)* and *lgg-1(G116AG117A)* mutants displayed a marked decrease of survival, *lgg-1(G116A)* mutants showed no difference with the wild-type animals. Moreover, the longevity of adults, which depends on bulk autophagy, was similar for *lgg-1(G116A)* and wildtype animals (Figure 2M) but strongly reduced for *lgg-1(G116AG117*)* mutants.

The autophagy capacity of LGG-1(G116A) protein, but not LGG-1(G116AG117*) or LGG-1(G116AG117A), was also documented by the elimination of apoptotic corpses in the embryo (Figure 2N, Supplementary FigS3) (Jenzer et al., 2019).

Overall, these data demonstrate that despite its defect to localize to autophagosome, LGG-1(G116A) achieves both selective and bulk autophagy during physiological and stress conditions. This is the first *in vivo* evidence that the autophagy functions of LGG-1/GABARAP can be uncoupled from its membrane conjugation. The non-functionality of LGG-1(G116AG117A) suggests that the precursor form is not responsible of LGG-1(G116A) autophagy activity. Despite an identical protein sequence, the truncated LGG-1(G116AG117*) is not functional for autophagy, indicating that the cleavage of the C-terminus from the precursor is essential for the functionality of LGG-1(G116A).

### The essential function of LGG-1 during development is unrelated to autophagy and independent of its cleavage and conjugation

The developmental phenotypes of the mutants *lgg-1(G116A), lgg-1(G116AG117A)* and *lgg-1(G116AG117*)* were then explored in embryo, larvae and adults and compared with *lgg-1(Δ)* and wild-type animals (Figure 3). We confirmed that *lgg-1(Δ)* homozygous animals present a massive lethality, during late embryogenesis or first larval stage (Figure 3B, H) (Manil-Ségalen et al., 2014). However, a careful analysis revealed that few escapers (circa 8% of the progeny), are able to reach adulthood and reproduce, allowing to maintain a *lgg-1(Δ)* homozygous population.

**Figure 3.**
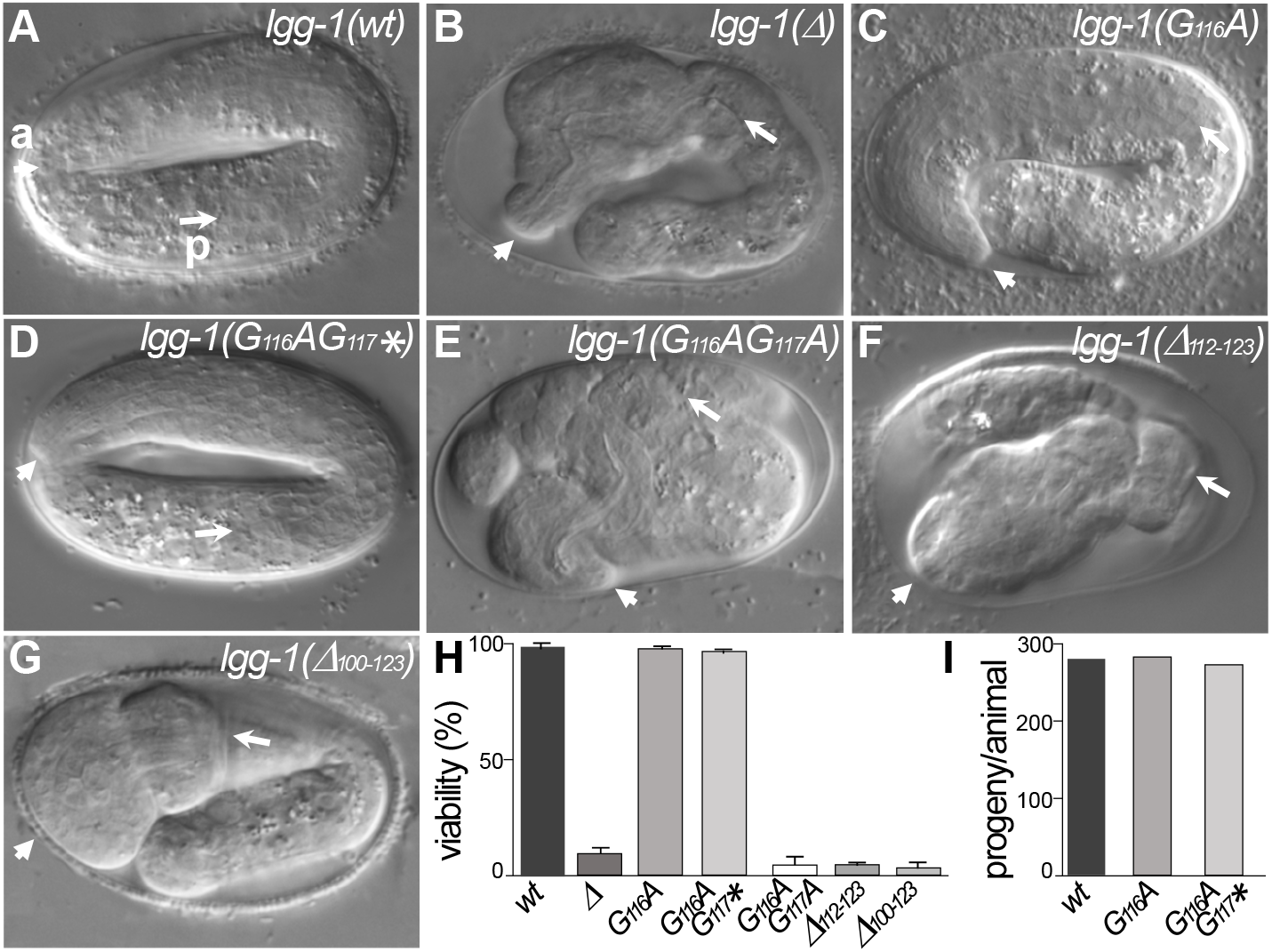
*lgg-1(G116A)* and *lgg-1(G116AG117*)* mutants are viable with no developmental defect. (**A-G**) DIC images of embryos after morphogenesis in *wild-type* (A), *lgg-1(Δ)* (B), *lgg-1(G116A)* (C), *lgg-1(G116AG117*)(D), lgg-1(G116AG117A)* (E), *lgg-1(Δ112-123)* (F), *lgg-1(Δ100-123)* (G). *lgg-1(G116AG117A), lgg-1(Δ112-123), lgg-1(Δ100-123), lgg-1(Δ)* mutant embryos present severe developmental defects. Short and long white arrows point to the anterior (a) and posterior (p) part of the pharynx, respectively. Scale bar is 10 μm. (**H**) The viability, expressed as the percentage of embryos reaching adulthood, is not affected in *lgg-1(G116A)* and *lgg-1(G116AG117*)* mutants (42<n< to 103). (**I**) The fertility, total number of progeny, of *lgg-1(G116A)* and *lgg-1(G116AG117*)* adults is similar to wild-type (n=20).

Neither *lgg-1(G116A)* nor *lgg-1(G116AG117*)* homozygous animals present any observable defect in development (Figure 3C, D, H) or adulthood and they reproduce at a similar rate than wild-type animals (Figure 3I). By contrast, *lgg-1(G116AG117A)* and the five independent mutants harboring various deletions and frameshifts of the C-terminus present a very strong lethality with the characteristic embryonic phenotype of *lgg-1(Δ)* animals (Figure 3E-H). Among them *lgg-1(Δ112-123)* presents a premature stop codon at position 112 and two others a frameshift in position 114 leading to an extension of the C-terminus (Figure 3 and Supplementary FigS1).

These data indicate that the cleaved LGG-1(G116A) and the truncated LGG-1(G116AG117*) forms, but not the precursor, are sufficient to recapitulate the normal development and viability, independently of membrane conjugation. The normal development of LGG-1(G116AG117*) demonstrates that the developmental functions of LGG-1 are independent of its autophagic functions. However, the C-terminus of the precursor is not compatible with LGG-1 activity for developmental processes.

Altogether, our data indicate that the cleavage of LGG-1 is necessary for all functions and suggest that autophagic but not developmental functions of LGG-1 form I require a further modification or activation.

### LGG-1(wt) and LGG-1(G116A) partially restore the survival to nitrogen starvation of yeast *atg8(Δ)*

Our results demonstrate that the localization of LGG-1 to the membrane is dispensable for autophagy and non-autophagy functions. To address whether it is a particularity of *C. elegans*, a similar strategy was performed for Atg8 in the yeast *S. cerevisiae*. Atg8 precursor ends with an arginine at position 117 (Supplementary FigS4A) and using mutant proteins expressed from centromeric plasmids the Oshumi lab (Kirisako et al., 2000; Nakatogawa et al., 2012) has shown that G116 is essential for autophagy. The endogenous *ATG8* was modified by an homologous recombination strategy to generate *atg8(G116A)* and *atg8(G116AR117*)* alleles, and the autophagy flux was assessed using the Pho8Δ60 reporter (Noda and Klionsky, 2008). Both *atg8(G116A)* and *atg8(G116AR117*)* mutants are unable to achieve a functional autophagy and behave similarly to *atg8(Δ)* or *atg1(Δ)* null mutants (Supplementary FigS4B). The analysis of nitrogen starvation survival showed that both *atg8(G116A)* and *atg8(G116AR117*)* strains were unable to recover after a 4 days starvation, similarly to *atg1Δ* and *atg8Δ* (Supplementary FigS4C).

The non-autophagy function of *atg8(G116A)* and *atg8(G116AR117*)* mutants was assessed by analyzing the shape of the vacuole (Banta et al., 1988). During exponential growth, *atg8(Δ)* cells frequently present multiple small vacuoles compared to wild-type cells which harbor usually less than 4 vacuoles (Supplementary FigS4D, E). The incidence of defective vacuolar shape decreases in *atg8(G116A)* and *atg8(G116AR117*)* cells indicating that the non autophagy functions are partially maintained. These data suggest that in *S. cerevisiae* the nonautophagy functions of Atg8 are partially independent of its cleavage and conjugation.

*Atg8* mutant was then used to investigate whether, the functionality of LGG-1(G116A) in autophagy is restricted to *C. elegans*. The capacity of LGG-1(G116A) to restore nitrogen starvation survival to *atg8(Δ)* mutant cells was compared with LGG-1(wt) and LGG-1(G116AG117*). The corresponding cDNA were cloned in a centromeric vector and the proteins were expressed in *atg8(Δ)* mutants. The expression of LGG-1(wt) and LGG-1(G116A), but not LGG-1(G116AG117*), improves the nitrogen starvation survival indicating a partial complementation (Supplementary FigS4F).This indicates that the functionality of LGG-1(G116A) is not restricted to *C. elegans*, supporting an intrinsic property of LGG-1 form I.

### Autophagy but not developmental functions of LGG-1(G116A) partially depends on LGG-2

Our previous study have shown a partial redundancy of LGG-1 and LGG-2 during starvation survival and longevity (Alberti et al., 2010), which raises the possibility of compensation of *lgg-1(G116A)* by the protein LGG-2. To analyze this possibility we used the large deletion mutant *lgg-2(tm5755)*, which is considered as a null (Manil-Ségalen et al., 2014), and constructed the double mutant strains *lgg-1(G116A);lgg-2(tm5755)* and *lgg-1(G116AG117*);lgg-2(tm5755)*.

Similarly to the single mutants *lgg-1(G116A)* and *lgg-2(tm5755)*, the double mutant *lgg-1(G116A);lgg-2(tm5755)* animals were viable and presented no morphological defect (Figure 4A-F). These data indicate that the correct development of *lgg-1(G116A)* is not due to a compensative mechanism involving *lgg-2*.

**Figure 4.**
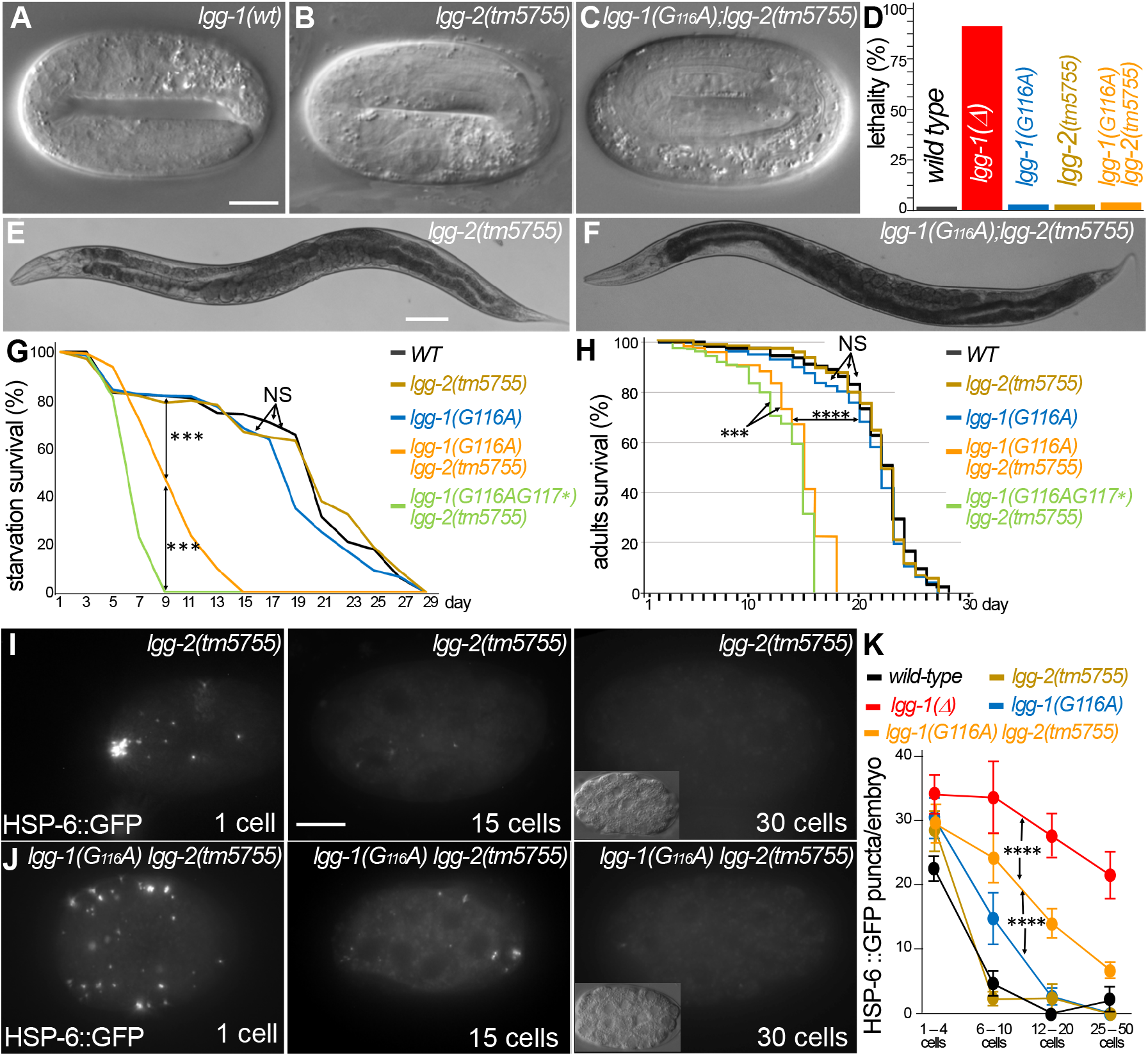
Autophagy but not developmental functions of LGG-1(G116A) partially depends on LGG-2. (**A-F**) DIC images of embryos and adults in *wild-type* (A), *lgg-2(tm5755)* (B, E), *lgg-1(G116A);lgg-2(tm5755)* (C, F). The double mutant *lgg-1(G116A);lgg-2(tm5755)* animals have no morphogenetic defects and no increase in lethality compare to single mutants or the *lgg-1(Δ)* (quantification in D). (**G-H**) Bulk autophagy during stress and aging was assessed by starvation survival (G, Chisquare test at day 9 ***p-value<0.001) and worm longevity (H, log rank test n>100 animals, ***p-value<0.001, ****p-value<0.0001). The survival of double mutants *lgg-1(G116A); lgg-2(tm5755)* and *lgg-1(G116AG117*); lgg-2(5755)* is reduced compared to *wild-type* and single mutant *lgg-1(G116A)* and *lgg-2(tm5755). lgg-1(G116A); lgg-2(tm5755)* animals survive to starvation better than *lgg-1(G116AG117*); lgg-2(5755)* and present a slightly higher lifespan. (**I-K**) *In vivo* epifluorescence imaging of paternal mitochondria (HSP-6::GFP) at the 1-cell, 15-cells and 30-cells stages in *lgg-2(tm5755)*, (I) *lgg-1(G116A);lgg-2(tm5755)* (J) embryos and quantification (n=50, 39, 35, 45, 46 Chi-square test ****<0.0001) (K). Elimination of mitochondria is efficient but delayed in *lgg-1(G116A);lgg-2(5755*) compared to *lgg-1(G116A)*. Insets show the corresponding DIC pictures. Scale bar is 10μm (A-C, I, J) or 100μm (D, E).

Next, we analyzed the autophagy functions in *lgg-1(G116A);lgg-2(tm5755)* animals. If LGG-2 compensates LGG-1(G116A) in autophagy, *lgg-1(G116A);lgg-2(tm5755)* animals should behave similarly to *lgg-1(G116AG117*);lgg-2(tm5755)* (of note *lgg-1(Δ);lgg-2(tm5755)* animals are not viable). The *lgg-1(G116A);lgg-2(tm5755)* animals presented a decrease for both survival to starvation and longevity compared to *lgg-1(G116A)* single mutant. However, they survived better than *lgg-1(G116AG117*);lgg-2(tm5755)* animals (Figure 4G, H). These results indicate that the functionality of LGG-1(G116A) in bulk autophagy partially relies on LGG-2. Selective autophagy during early embryogenesis was then quantitatively analyzed in the double mutant strains (Figure 4I-K). Surprisingly, paternal mitochondria were eliminated in the *lgg-1(G116A);lgg-2(tm5755)* animals indicating that LGG-1(G116A) is sufficient for the allophagy process. It suggests that paternal mitochondria could be degraded by autophagosomes devoid of both LGG-1 and LGG-2. However, a delay in the degradation was observed compared to *lgg-1(G116A)* animals suggesting that the autophagy flux is reduced. These results reveal a partial redundancy between LGG-1 and LGG-2 in autophagy, but demonstrate that LGG-1(G116A) fulfills the developmental function and maintains some autophagy activity independently of LGG-2.

Interestingly, this detailed analysis also revealed a slight delay in the elimination of paternal mitochondria in *lgg-1(G116A)* animals compared to wild-type (Figure 4K). Although the cleaved LGG-1 is sufficient for autophagy, this observation suggests that the loss of membrane targeting could affect the dynamic of autophagy flux.

### The degradation of autophagosome is delayed in LGG-1(G116A)

The observation that LGG-1(G116A) degrades paternal mitochondria slower led us to investigate whether a particular step of the autophagy flux could be modified in the absence of lipidation. The autophagic flux and the dynamic of autophagosome formation were compared between *lgg-1(G116A), lgg-1(G116AG117*)* and *lgg-1(Δ)* animals. We first focused on the early embryo where the autophagy process is stereotyped and the nature of the cargos and the timing of degradation well characterized. Moreover, the autophagosomes sequestering the paternal mitochondria are clustered and positive for LGG-2 (Figure 5A)(Manil-Ségalen et al., 2014). In *lgg-1(Δ)* mutant, LGG-2 autophagosomes are not detected as a cluster but are spread out in the whole embryo as single puncta, which persist after the 15-cells stage (Figure 5B, E). This indicates that individual LGG-2 structures can be formed in absence of LGG-1, but are not correctly localized and not degraded properly, presumably because of the role of LGG-1 in cargo recognition (Sato et al., 2018) and of a latter involvement in the maturation of autophagosomes, respectively. The pattern of LGG-2 is somehow different in *lgg-1(G116A)* and *lgg-1(G116AG117*)* mutants, forming sparse structures of heterogeneous size, which persist longer (Figure 5C-F). These data suggest that the cleaved and the truncated LGG-1 can both promote the recruitment of LGG-2 to autophagic structures but display an altered autophagic flux. The analysis of the colocalization between paternal mitochondria and LGG-2 did not reveal an increase in *lgg-1(G116A)* or *lgg-1(G116AG117*)* mutants (Figure 5G-J and Supplementary FigS5). These data suggest that the elimination of paternal mitochondria in *lgg-1(G116A)* animals is not due to the enhanced recruitment of LGG-2. A western blot analysis of adult worm indicates that there is no increase of LGG-2 expression in *lgg-1(Δ), lgg-1(G116A)* and *lgg-1(G116AG117*)* mutants (Figure 5K). The absence of enhanced colocalization between LGG-2 and paternal mitochondria suggests that in *lgg-1(G116A)* the paternal mitochondria are degraded by autophagosomes devoid of lipidated LGG-1 and LGG-2.

**Figure 5.**
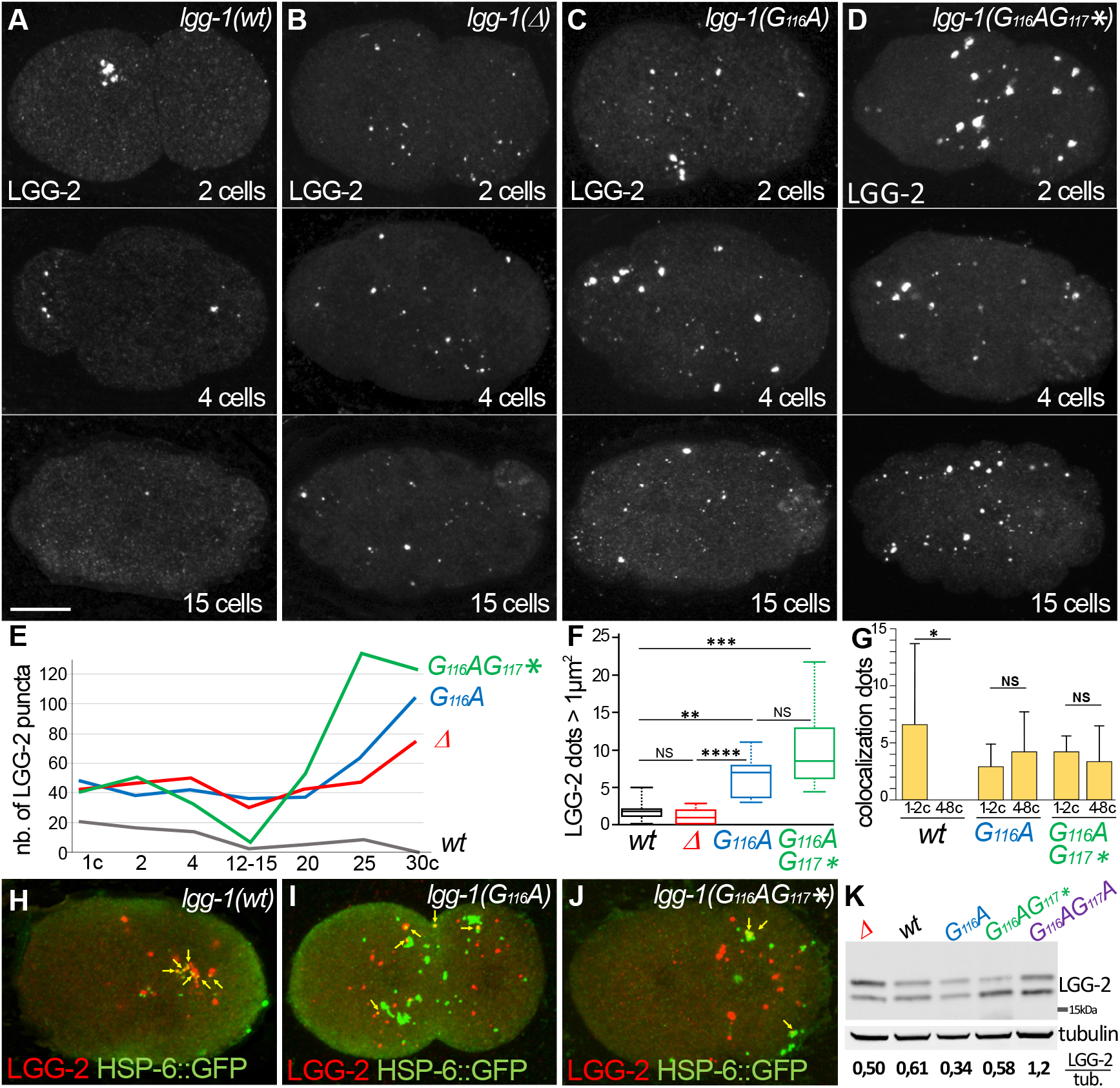
The degradation of autophagosome is delayed in LGG-1(G116A) (**A-F**) Confocal images of LGG-2 immunofluorescence in 2 cells, 4 cells and 15 cells in *wild-type* (A), *lgg-1(Δ)* (B), *lgg-1(G116A)* (C), *lgg-1(G116AG117*)* (D) and quantification of the number (E) and size of puncta (F)(embryo analyzed 19, 37, 28, 14; Mann-Whitney test, p-value ****<0.0001). In *lgg-1(G116A)* and *lgg-1(G116AG117*)* mutants LGG-2 is detected as heterogeneous sparse structures that persist. (**G-J**) Colocalization analysis of paternal mitochondria (HSP-6::GFP) and LGG-2 puncta (H) from confocal images of *wild-type* (H), *lgg-1(G116A)* (I) and *lgg-1(G116AG117*)* (J) early embryos. (Mean + SD, n= 16, 20, 12, Kruskal Wallis test p-value*<0.05). The clustering of paternal mitochondria and LGG-2 autophagosomes is absent in *lgg-1(G116A)* and *lgg-1(G116AG117*)* where HSP-6::GFP and LGG-2 puncta are mainly separated with rare colocalization events (yellow arrows). (**K**) Western blot analysis of endogenous LGG-2 from total protein extracts from *wild-type, lgg-1(G116A), lgg-1(G116AG117*), lgg-1(G116AG117A), lgg-1(Δ)* young adults. The quantification of total LGG-2 was normalized using tubulin. *(See also supplementary Figure S5)*.

The further characterization of autophagic structures in *lgg-1(G116A)* and *lgg-1(G116AG117*)* embryos was performed by electron microscopy and compared with wild-type and *lgg-1(Δ)* mutant embryos (Figure 6). In wild-type animals, autophagosomes containing cytoplasmic materials (referred as type 1 Figure 6A, H) and the characteristic paternal mitochondria (Zhou et al., 2016) were observed in 1-4 cells embryos, and less frequently in 5-10 cells embryos. At that stage, rare autophagosomes containing partially degraded material are present (referred as type 2). As expected, almost no autophagosome was observed in *lgg-1(Δ)* embryos and paternal mitochondria were non-sequestered (Figure 6B, H). In *lgg-1(G116A)* embryos, the numbers of type 1 and type 2 autophagosomal structures increase both in 1-4 cells and 5-10 cells stages. The autophagosomes look closed and contain various cellular materials and membrane compartments (Figure 6C-E). This confirms that LGG-1(G116A) is sufficient to form functional autophagosomes but with a delayed degradation. On the other hand, *lgg-1(G116AG117*)* embryos present non-sequestered paternal mitochondria (Figure 6F) and multi-lamellar structures containing cytoplasm but devoid of membrane organelles (type 3 Figure 6G, H). Type 3 structures are not seen in wild-type, *lgg-1(Δ)* and *lgg-1(G116A)*, suggesting a gain of function of the truncated LGG-1(G116AG117*) protein that induces a non functional compartment.

**Figure 6.**
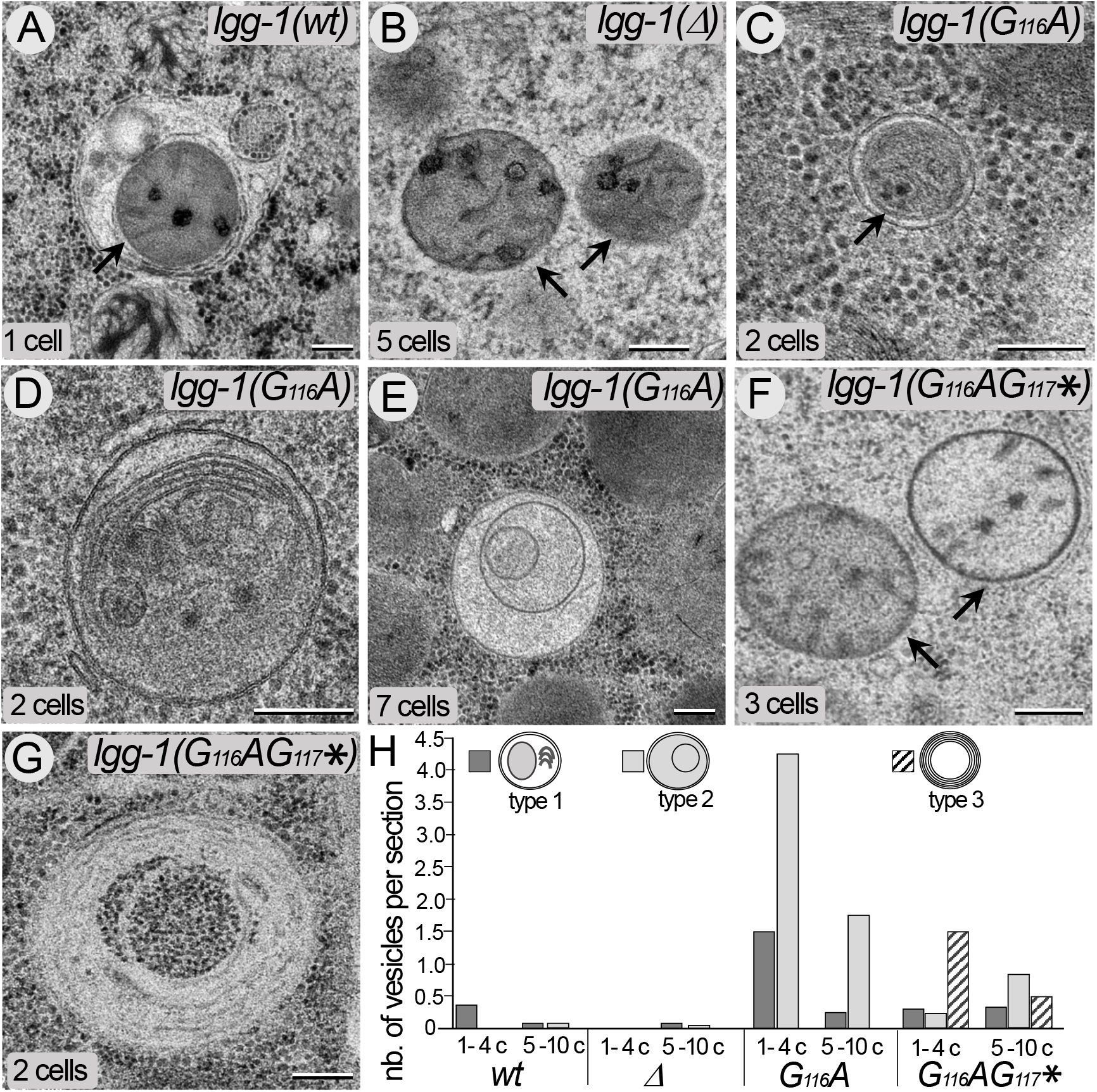
The cleaved but not the truncated LGG-1 is sufficient for autophagosome biogenesis. (**A-G**) Electron microscopy images of autophagosomes in *wild-type* (A), *lgg-1(Δ)* (B), *lgg-1(G116A)* (C-E) and *lgg-1(G116AG117*)* (F-G) early embryos. Type 1 autophagosomes appear as closed structures containing various membrane organelles. Among those, sequestered paternal mitochondria (black arrows) are observed in *wild-type* and *lgg-1(G116A)* embryos but remain unsequestered in *lgg-1(Δ)* and *lgg-1(G116AG117*)* embryos. Type 2 autophagosomes appear as closed structures containing unidentified or degraded materials. Type 3 structures (G) are multi-lamellar structures only detected in *lgg-1(G116AG117*)* embryos. Scale bar is 200 nm. (**H**) Quantification of type1, type 2 and type 3 structures at 1-4 cells and 5-10 cells stages. In *lgg-1(G116A)* embryos, the numbers of type 1 and type 2 autophagosomal structures increase supporting a retarded degradation. (n sections= 19, 13, 14, 48, 16, 16, 13, 6, n embryos= 7, 5, 7, 13, 4, 4, 9, 2).

Altogether, these data indicate that the cleaved, but not the truncated, LGG-1 form I is able to form functional autophagosomes but with a delayed degradation.

### The cleaved LGG-1 is sufficient for UNC-51/ULK1 dependent autophagosome initiation and biogenesis

Next, we analyzed another physiological autophagy process to further characterize the initiation and elongation of autophagosomes in *lgg-1(G116A)* and *lgg-1(G116AG117*)* mutants (Figure 7A-D). Using powerful genetic approaches the Zhang lab has previously described the aggrephagy pathway in *C. elegans* embryo (Lu et al., 2011; Tian et al., 2010; Wu et al., 2015; Zhang et al., 2009). In the 50-100 cells embryo we quantified the colocalization between ATG-18/WIPI2 and LGG-2. ATG-18, the worm homolog of the omegasome marker WIPI2 (Polson et al., 2010), acts at an early step of biogenesis (Lu et al., 2011). Puncta labelled with ATG-18 only, both ATG-18 and LGG-2, or LGG-2 only were considered as omegasomes, phagophores and autophagosomes, respectively. In *lgg-1(RNAi)* animals the number of omegasomes increases while the proportion of phagophore decreases compared to the wildtype embryos (Figure 7A, B, E, F). This indicates that the initiation of autophagy is triggered in absence of LGG-1, but the extension of autophagosome is defective. *lgg-1(G116A)* animals show no difference with the wild-type (Figure 7C, E-G) supporting that both initiation and phagophore extension are normal with the cleaved LGG-1. Similarly to the *lgg-1(RNAi), lgg-1(G116AG117*)* animals are defective in the phagophore extension (Figure 7D-F). These data confirm that the cleaved LGG-1(G116A), but not the truncated LGG-1(G116AG117*), is fully functional for the early step of autophagosome biogenesis.

**Figure 7.**
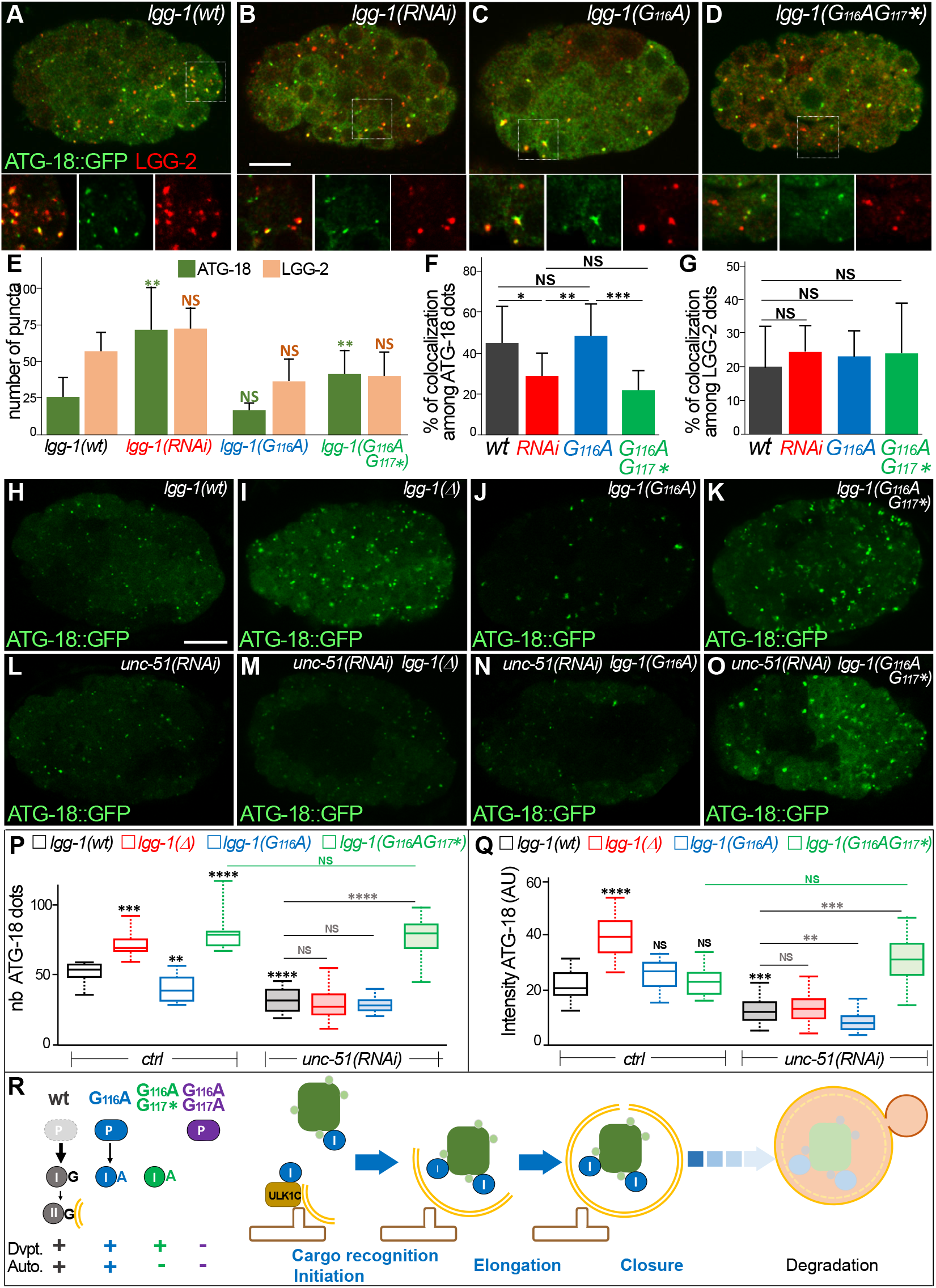
The cleaved LGG-1 is sufficient for UNC-51/ULK1 dependent autophagosome initiation and biogenesis. (**A-D**) Confocal images of ATG-18::GFP (green) and LGG-2 (red) immunofluorescence in *wildtype* (A), *lgg-1(RNAi)* (B), *lgg-1(G116A)* (C), *lgg-1(G116AG117*)* (D) embryos. Insets are 2-fold magnification of the white boxed regions. Scale bar is 10μm. (**E-G**) Quantification of ATG-18 and LGG-2 puncta (E) and colocalization events (F, G) (mean + SD, n= 10, 10, 10, 10; Kruskal Wallis p-value*<0.05**<0.01***<0.001). A decreased colocalization is observed in *lgg-1(RNAi)* and *lgg-1(G116AG117*)* but not *lgg-1(G116A)* embryos. (**H-O**) *In vivo* confocal images of ATG-18::GFP, showing the initiation of autophagosome biogenesis, in 50-100 cells embryos for *wild-type* (H), *lgg-1(Δ)(I), lgg-1(G116A)* (J), *lgg-1(G116AG117*)* (K), *unc-51 (RNAI)(L), unc-51(RNAI);lgg-1(tm3489)(M), unc-51(RNAI);lgg-1(G116A)(N)* and *unc-51(RNAI);lgg-1(G116AG117*)(O)*. Scale bar is 10μm. (**P-Q**) Boxplots of the number of ATG-18::GFP puncta (P) (n=13, 24, 10, 14, 28, 36, 27,20) and the mean fluorescent intensity (Q)(n= 42, 45, 55, 140, 145, 130,115, 100) (Kruskal-Wallis test p-value *<0.05, **p<0.01 ***p<0.001, ****p<0.0001, NS non-significant) **(R)** Model of the results presented in this study. The precursor form of LGG-1 is not able to promote the autophagy (Auto.) and developmental functions (Dvpt.). Both the cleaved and the truncated form I are sufficient for development but the truncated is not sufficient for development. LGG-1(G116A) is not lipidated but sufficient for initiation and autophagosome biogenesis. A delay in autophagosome degradation (discontinuous blue arrow) suggests that the conjugation to autophagosome membrane is important for the late stages of autophagy.

Data in yeast and mammals have revealed that Atg8/LC3/GABARAP can interact with Atg1/ULK1 and modify the kinase activity of the ULK1 complex (Alemu et al., 2012; Grunwald et al., 2020; Kraft et al., 2012; Nakatogawa et al., 2012). In *C. elegans*, LGG-1 can directly binds UNC-51/ULK1 (Wu et al., 2015) and the cargo SEPA-1 (Zhang et al., 2009) and could have an early function for initiating aggrephagy (Lu et al., 2011). To decipher the function of LGG-1 form I in the induction of autophagy we performed a genetic approach. Using RNAi we depleted UNC-51 and quantified the initiation events *in vivo* in wild-type, *lgg-1(Δ), lgg-1(G116A)*, and *lgg-1(G116AG117*)* embryos (Figure 7H-Q). As expected, depleting UNC-51 results in the decrease of ATG-18 puncta while depleting LGG-1 leads to the increase of ATG-18 intensity and number of puncta, indicative of its downstream functions. The decrease of ATG-18::GFP puncta after co-depletion of UNC-51 and LGG-1 supports an earlier function for UNC-51 than for LGG-1. For *lgg-1(G116A)* animals, a small decrease was observed in the number of puncta compared to the wild-type animals but no change in the total signal of ATG-18::GFP. Moreover, the depletion of UNC-51 further decreased ATG-18::GFP puncta and intensity confirming that LGG-1(G116A) almost behaves like the wild-type LGG-1 for the initiation (Figure 7J, N, P-Q).

In *lgg-1(G116AG117*)* embryos, a marked increase of ATG-18 puncta was observed, but contrarily to *lgg-1(Δ)*, the number did not decrease when UNC-51 was depleted (Figure 7K, O-Q). This data confirms electron microscopy observations and supports a gain of function for the truncated LGG-1(G116AG117*), independent of ULK1 complex.

Altogether, the analyses of LGG-1(G116A) indicate that many of the functions of LGG-1 in autophagy can be achieved by the cleaved but non lipidated form I, including cargo recognition, initiation and autophagosome biogenesis (Figure 7R). However, the lipidation of LGG-1 appears to be important for the final steps of maturation/degradation of the autophagosome.

## Discussion

The most surprising result of this study is the discovery that LGG-1(G116A) is functional for all the autophagy processes that have been tested, covering physiological or stress conditions and selective or bulk autophagy. To our knowledge, it is the first report demonstrating that different autophagy processes are fully achieved *in vivo* in a non lipidable mutant. Addressing the similar question in mammals is challenging. In cultured cells, an elegant CRISPR strategy allowed to knock out together the six LC3/GABARAP homologs, but point mutations of the conserved glycine have not been reported (Nguyen et al., 2016). Most of the studies on the terminal glycine used transgenic overexpression constructs (Chen et al., 2007; Kabeya et al., 2004). Interestingly, one study reported that part of the autophagy functions of GABARAPL1 is independent of its lipidation (Poillet-Perez et al., 2017). Several studies have used mutations in the conjugation machinery (Atg3, Atg5, Atg7) or the Atg4 protease to analyze the role of the form I, respectively (Hill et al., 2019; Hirata et al., 2017; Nishida et al., 2009; Ohnstad et al., 2020; Vujić et al., 2021). An alternative autophagy has been reported in mutants of the conjugation machinery (Nishida et al., 2009), but blocking the conjugation system presumably affects all LC3/GABARAP homologs. Moreover, the presence of four homologs of Atg4 in mammals, which specificity versus LC3/GABARAP is unknown, and the dual role in the cleavage of the precursor and the delipidation entangle the analysis of the phenotypes.

Our data show no evidence for an intrinsic function of the LGG-1 precursor but the importance of its active cleavage. It is not unexpected because in many species the Atg8 precursor is not detected, suggesting that the cleavage occurs very soon after or even during translation. Moreover, phylogenetic analyses of LC3/GABARAP show no conservation in sequence and length of the C-terminus but the presence of at least one residue after the conserved G116. The hypothesis of a selective constraint on the cleavage but not on the C-terminus sequence could explain the persistence of a precursor form. Further studies would be necessary to address the precise implication of the di-glycine G116G117 in the process.

Albeit a similar sequence, the difference of functionality of the cleaved LGG-1(G116A) and the truncated LGG-1(G116AG117*) suggests that the cleavage allows a first level of specificity for LGG-1 functions. Our data is the first evidence that LGG-1 function in development relies on the cleavage but is independent of autophagy and conjugation. Our results could explain the embryonic lethality reported upon depletion of the two Atg4 homologs precursors in *C. elegans* (Wu et al., 2012). While the cleavage is sufficient for the developmental functions, the autophagy functions of LGG-1 form I seem to require a further modification to be active. Our data suggest that this modification is dependent and possibly associated to the cleavage.

Our observations in yeast also support an autophagy independent function of Atg8 form I in vacuolar shaping. Non autophagic functions for LC3/GABARAP have been identified in yeast and higher eukaryotes (Ishii et al., 2019; Liu et al., 2018; Schaaf et al., 2016; Wesch et al., 2020), but the roles of the cytosolic forms are poorly documented especially in the context of the development. The two Atg8 homologs of drosophila are involved in several developmental processes independently of canonical lipidation (Chang et al., 2013) or autophagy (Jipa et al., 2020). They are highly similar and both correspond to GABARAP homologs (Manil-Ségalen et al., 2014). It is possible that duplication of Atg8 during evolution allowed the acquisition of specific developmental functions by GABARAP proteins but reports in apicomplex parasites (Lévêque et al., 2015; Mizushima and Sahani, 2014) rather supports a non autophagy ancestral function of Atg8.

A goal of this study was to bring new insights concerning the implication of LGG-1 form I in various steps of autophagy. Numerous studies identified interacting partners of Atg8/LC3/GABARAP family during autophagy but its mechanistic function for autophagosome biogenesis is still debated. In yeast, the amount of Atg8 regulates the level of autophagy, controls phagophore expansion but is mainly released from the phagophore assembly site during autophagosome formation (Xie et al., 2008). *In vitro* studies using liposomes or nanodiscs suggested that Atg8 is a membrane-tethering factor and promote hemifusion (Nakatogawa et al., 2007), membrane tubulation (Knorr et al., 2014) or membrane-area expansion and fragmentation (Maruyama et al., 2021). Another study shows that Atg8–PE assembles with Atg12–Atg5-Atg16 into a membrane scaffold that is recycled by Atg4 (Kaufmann et al., 2014). A similar approach with LGG-1 supports a role in tethering and fusion activity (Wu et al., 2015). *In vivo*, the functions of these proteins presumably depend of their amount, their posttranslational modifications and the local composition of the membrane. For instance, an excess of lipidation of the overexpressed LGG-1 form I mediates the formation of enlarged protein aggregates and impedes the degradation process (Wu et al., 2015). A recent report showed that the phosphorylation of LC3C and GABARAP-L2 impedes their binding to ATG4 and influences their conjugation and de-conjugation (Herhaus et al., 2020). Our genetic data indicate that form I of LGG-1 is sufficient for initiation, elongation and closure of autophagosomes but that lipidated LGG-1 is important for the dynamic of degradation. However, the partial redundancy with LGG-2 is presumably a factor important for the late maturation and degradation process.

The mechanism allowing the effective degradation of specific cargoes in *lgg-1(G116A)* mutant is unclear but one can assume that physical interactions, through LIR or other domains, are sufficient to initiate the extension of the phagophore around the cargoes. If the main functions of LGG-1 reside in its capacity to bind multiple proteins, the localization to autophagosome membrane through lipidation is an efficient but not unique way to gather cargoes and autophagy complexes.

Finally, the observation that paternal mitochondria could be degraded by autophagosomes devoid of both LGG-1 and LGG-2 is intriguing and further exemplifies the high level of plasticity and robustness of autophagy.

## Acknowledgments

The authors would like to thank Thorsten Hoppe for providing LGG-1 antibodies, Fulvio Reggiori for yeast plasmids, and the Caenorhabditis Genetic Center, which is funded by the NIH National Center for Research Resources (NCRR), for strains. We are grateful to Laïla Sago and Virginie Redeker for help with mass spectrometry. The present work has benefited from the facilities and expertise of the I2BC proteomic platform (Proteomic-Gif, SICaPS) supported by IBiSA, Ile de France Region, Plan Cancer, CNRS and Paris-Sud University as well as the core facilities of Imagerie-Gif, member of IBiSA), supported by “France-BioImaging” and the Labex “Saclay Plant Science”.

This work was supported by the Agence Nationale de la Recherche (project EAT, ANR-12-BSV2-018), the Association pour la Recherche contre le Cancer (SFI20111203826) and the Ligue contre le Cancer (MM). RoL received a fellowship from Fondation pour la Recherche Médicale.

## Author contributions

Conceptualization, Re.L. and E.C.; Methodology, Ro.L., Ch.L., and E.C.; Investigation, Ro.L., Ce.L., Ch.L., G.Q., M.P., M.H.C. and E.C.; Formal analysis, Ro.L., Ce.L., M.P., and M.H.C.; Validation, Ro.L., Ce.L., M.P., M.H.C. and R. L.; Visualization, Ro.L., Ce.L., M.P., M.H.C., and Re.L.; Writing – Original Draft, Re.L.; Writing –Review & Editing, Ro.L., M.H.C., E.C. and Re.L.; Funding Acquisition, Re.L.; Resources, M.R. and G.Q.; Supervision, Ch.L., E.C., M.H.C. and Re.L.; Project administration, Re. L.

## Supplemental figures titles and legends

**Supplementary Figure S1.**
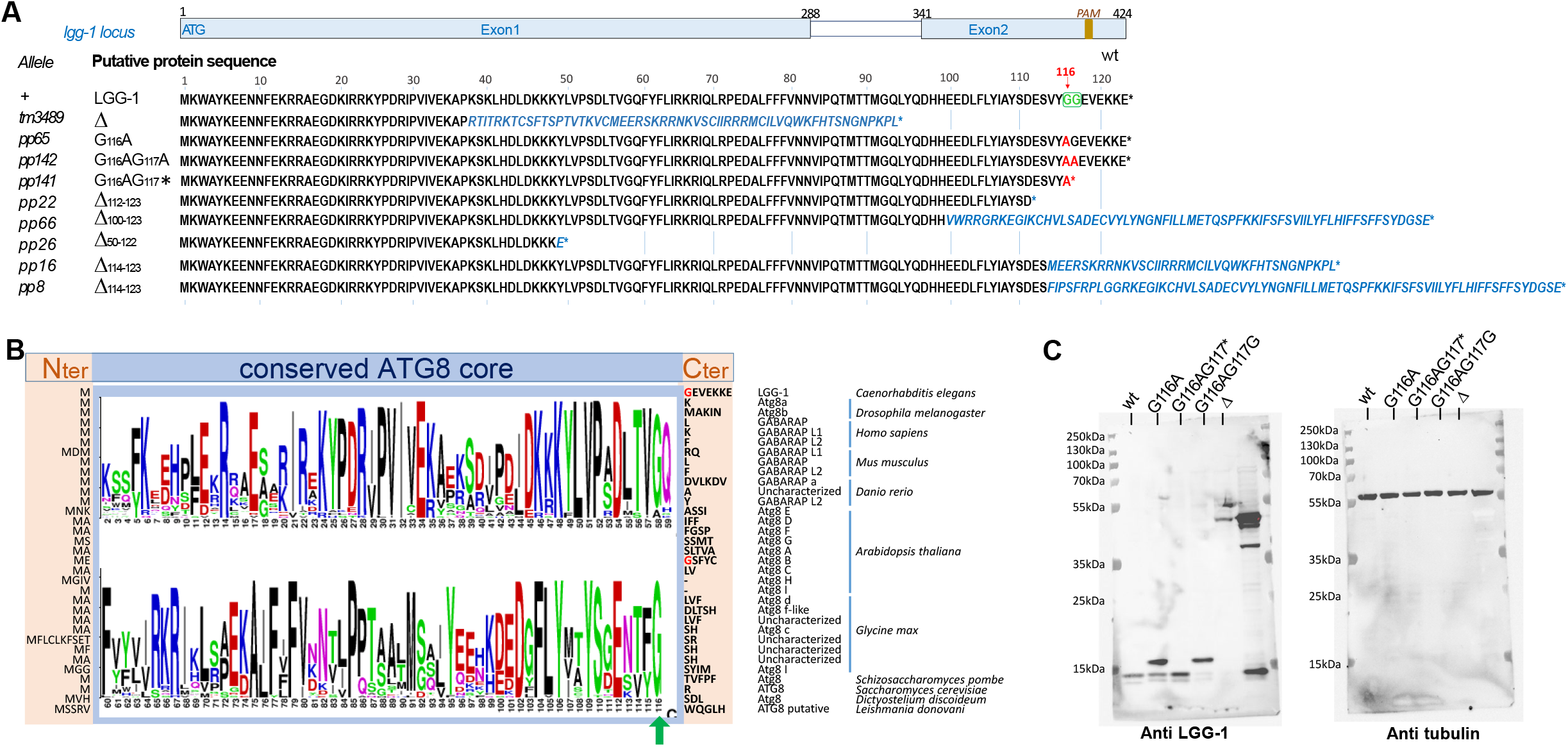
(related to Figure 1) - Description of *lgg-1* alleles. **(A**) Schematic representation of *lgg-1* locus indicating the exon/intron composition and the localization of the PAM used for CRISPR-Cas9. The names of the alleles and the theoretical sequences of the proteins series are indicated. Residues in red are the results of the point mutations and residues in blue italics are the theoretical consequence of frameshifts. The diglycine motif in position 116-117 is shown in green and the red arrow points to the conserved glycine 116. **(B)** Comparison of Atg8/GABARAP homologs in worm (Ce), fly (Dm), human (Hs), mouse (Mm), zebrafish (Dr), plants (At, Gm), yeasts (Sp, Sc), amoeba (Dd) and parasite protozoa (Ld), identified in the eukaryotic proteomes from NCBI landmark Blast database. The conserved core sequence (blue box) is represented using a LOGO analysis (after removal of a small insertion in At Atg8D and in two Gm uncharacterized homologs) and the variable N- and C-terminus sequences are indicated in light beige. Green arrows points to the invariable glycine 116. (**C**) Uncropped western blots shown in Figure 1C merged with molecular weight markers. Right sample is unrelated to this work.

**Supplementary Figure S2.**
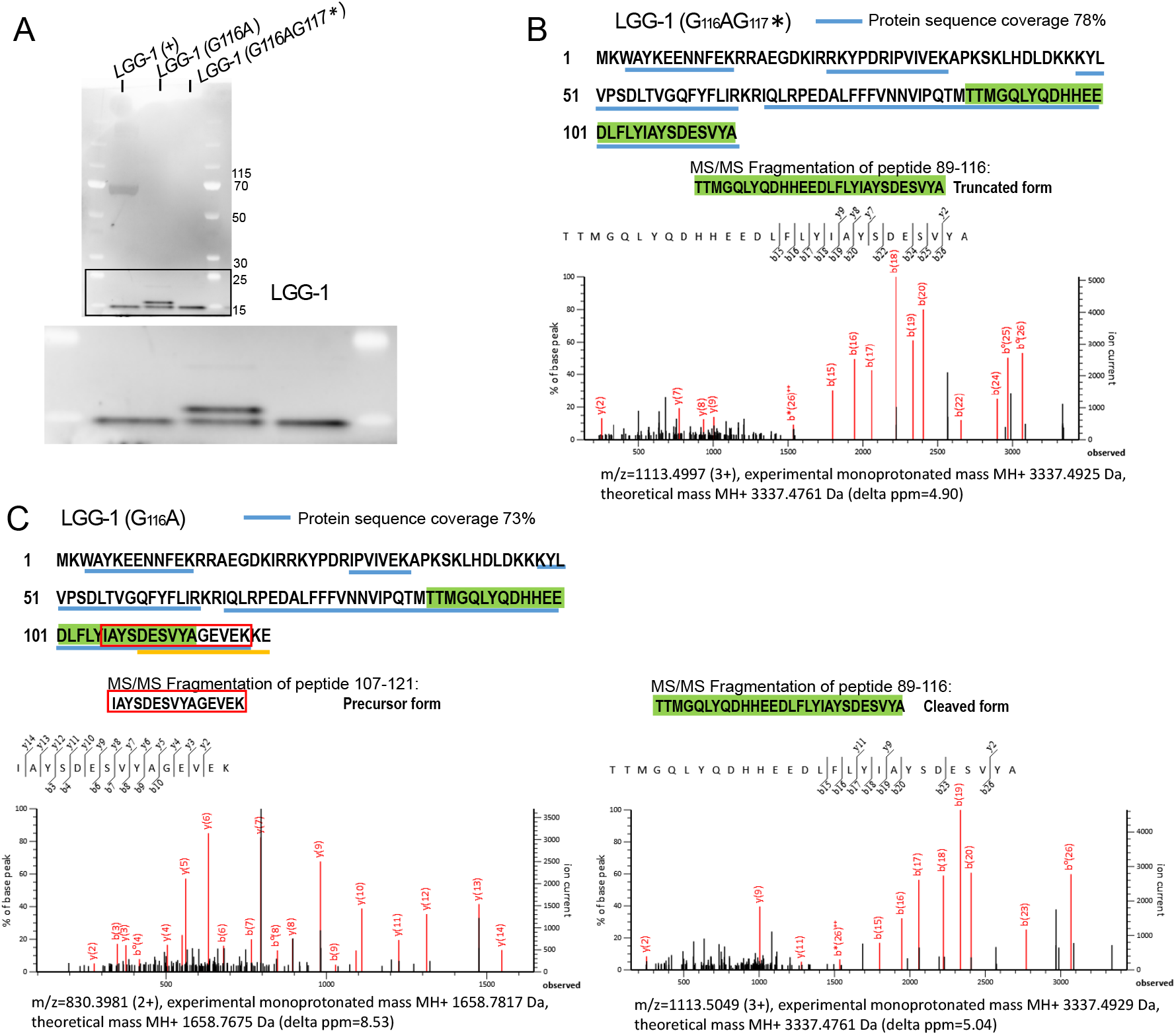
(related to Figure 1) - Identification of LGG-1(G116A) and LGG-1(G116AG117*) forms. A) Western blot analysis of affinity purified samples using LC3 traps. Molecular weight markers (kDa) are indicated on the right. B) Protein sequence, and peptides coverage of LGG-1(G116AG117*) identified by mass spectrometry analyses after trypsin treatment (blue underline). The MS/MS fragmentation of the C-terminal peptide 89-116 (green) identifying the truncated form is shown below. C) Protein sequence and peptides coverage of LGG-1(G116A) identified by mass spectrometry analyses after trypsin treatment (blue underline). The yellow underline indicates a C-terminal 111-123 peptide identified after trypsin and AspN digestion. The MS/MS fragmentation of the C-terminal peptide 107-121 (red box) identifying the precursor form and peptide 89-116 (green) identifying the cleaved form are shown below.

**Supplementary Figure S3.**
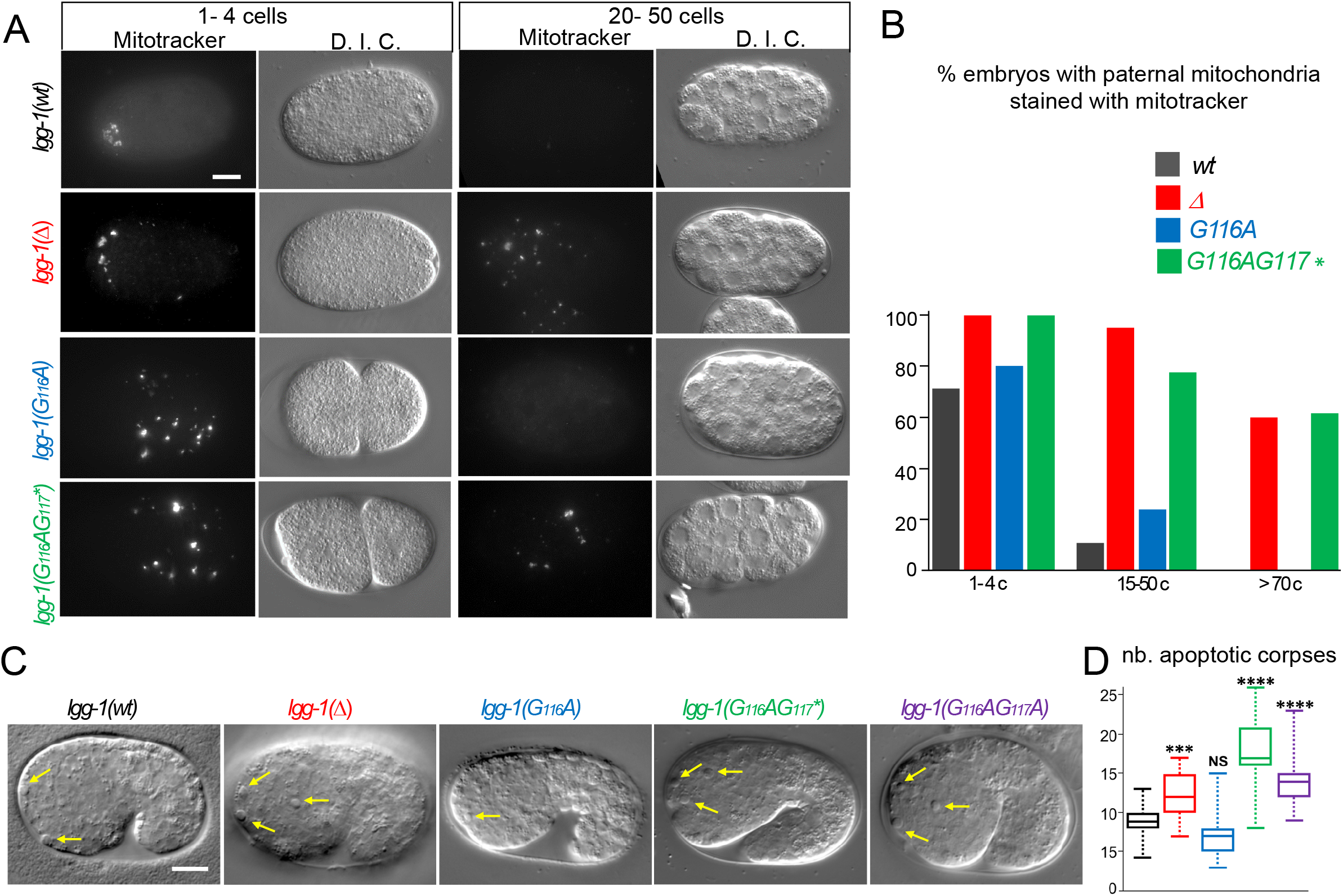
(related to Figure 2) - Autophagy is functional in *lgg-1(G116A)* but not in *lgg-1(G116AG117*)* (**A-B**) *In vivo* epifluorescence images of paternal mitochondria (left, mitotracker) and corresponding DIC images (right) at the 1-4 cells and 20-50 cells stages in *wild-type, lgg-1(Δ), lgg-1(G116A), lgg-1(G116AG117*)* embryos showing an effective degradation of paternal mitochondria in *wt* and *lgg-1(G116A)* but not in *lgg-1(Δ) lgg-1(G116AG117*)*. Quantification are shown in (B). (**C-D**) DIC images of 1.5 fold stage *wild-type, lgg-1(Δ), lgg-1(G116A), lgg-1(G116AG117*)* and *lgg-1(G116AG117A)* embryos. Apoptotic corpses in the head region are indicated by yellow arrows. The box-plots quantification in D is the same shown in Figure 2N. Scale bar is 10 μm.

**Supplementary Figure S4.**
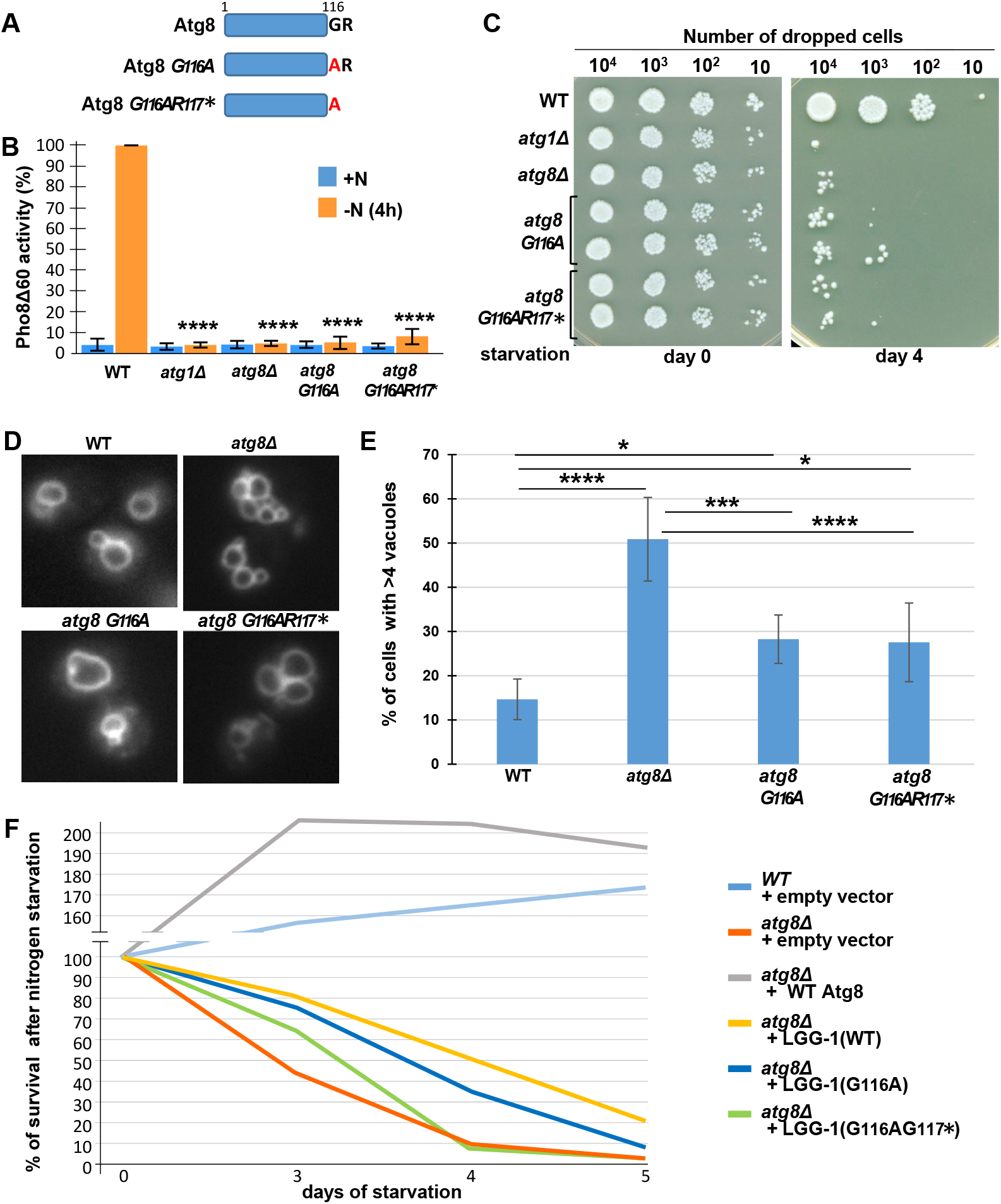
Atg8(G116A) and Atg8(G116AR117*) are functional for vacuolar shaping but not for autophagy in *S. cerevisiae*. **(A)** Schematic representation of wild-type and mutant Atg8 proteins. **(B)** Autophagic activity is abolished in *S. cerevisiae* mutant strains *atg8G116A* or *atg8G116A-R117Δ* forms of Atg8. Autophagic activity of wild-type, *atg1Δ, atg8Δ, atg8G116A* and *atg8G116AR117\** strains was measured using the quantitative Pho8Δ60 assay, before (+N) and after (-N) 4 hours of nitrogen starvation. The data shown are the mean ± s.e.m. of 3 independent experiments (Mann-Whitney test, p-value****<0.0001). **(C)** S. *cerevisiae* mutants *atg8Δ, atg8G116A* or *atg8G116AR117\** have a similar cell viability defect upon nitrogen starvation. **(D)** Epifluorescence image of FM4-64 staining of the vacuole of *wild-type* and *atg8* mutant strains (single focal plan). **(E)** Quantification of vacuoles in wild-type and *atg8* mutants*. atg8G116A* and *atg8G116AR117\** mutants present a less severe vacuolar phenotype than *atg8Δ* mutant (n > 200 cells, Chi-square test, p-value *<0.05, ***<0.001, ****<0.0001). **(F)** Rescue assays of *atg8Δ* by *lgg-1(wt), lgg-1(G116A)* and *lgg-1(G116AG117*)*. One representative of three independent experiments is shown. The survival in nitrogen starvation of *atg8Δ* cells transformed with *lgg-1(wt), lgg-1(G116A), lgg-1(G116AG117*)* or *ATG8* was compared with *atg8Δ and wild-type*. The percentage of surviving cells was calculated in comparison with day 0. *lgg-1(G116A)* expression improves the survival but less efficiently than *lgg-1(wt)*.

**Supplementary Figure S5.**
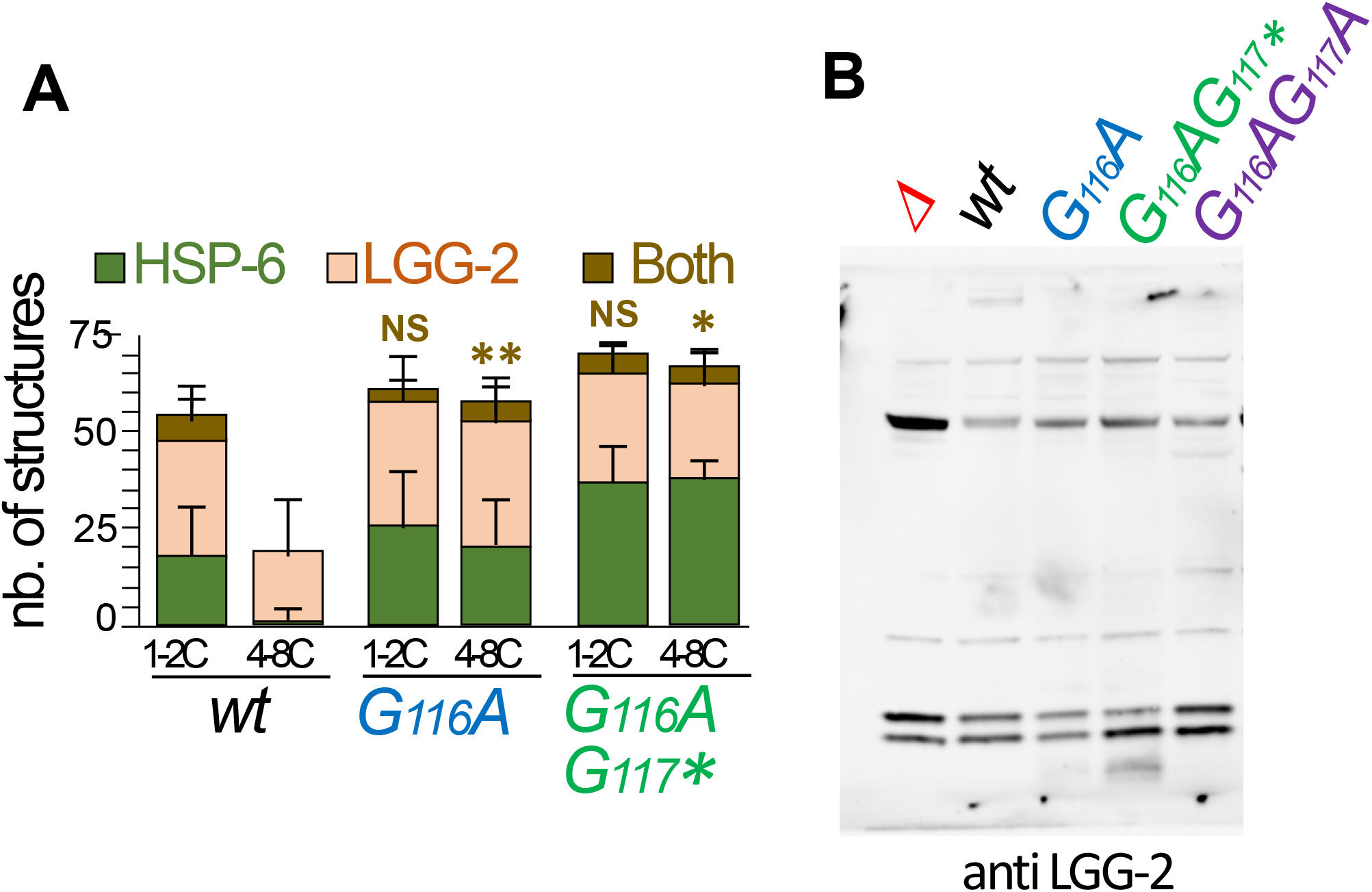
(related to Figure 5) – (**A**) Quantification of puncta positive for the paternal mitochondria marker HSP-6::GFP, the autophagosomal marker LGG-2 and both in early embryos *wild-type, lgg-1(G116A)* and *lgg-1(G116AG117*)* (Mean + SD, n= 16, 20, 12 Kruskal Wallis p-value*<0.05, **<0.01)(related to figure 5G). (**B**) Uncropped image of the western blot shown in Figure 5K.

## RESOURCE AVAILABILITY

Further information and requests for resources and reagents should be directed to the corresponding author, Renaud Legouis.

## EXPERIMENTAL MODEL AND SUBJECT DETAILS

### *C. elegans* culture and strains

Nematode strains were grown on nematode growth media (autoclaved 1.5g NaCl (Sigma-Aldrich, 60142), 1.5g bactopeptone (Becton-Dickinson, 211677), 0.5ml cholesterol (Sigma-Aldrich, C8667) 5mg/ml, 10g bacto agar (Becton-Dickinson, 214010) supplemented with 500μl CaCl2 (Sigma-Aldrich, C3306) 1M, 500μl MgSO4 (Sigma-Aldrich, M5921) 1M, 10ml KH2PO4 (Sigma-Aldrich, P5655) 1M, 1650μl K2HPO4 (Sigma-Aldrich, 60356) 1M and fed with *Escherichia coli* strain OP50. The *C. elegans* Bristol N2 strain was used as a wild-type strain. Other strains and genotypes are listed in Supplementary Table 1.

### *S. cerevisiae* culture and strains

Yeast cells were grown to log phase in YPD (1% yeast extract, 2% bactopeptone and 2% glucose) or complete synthetic medium (CSM) without uracil or leucine. The reference strain is BY4742. Other strain and genotypes are listed in Supplementary Table 1.

## METHOD DETAILS

### CRISPR-Cas9

We used CRISPR-Cas9 approach optimized for *C. elegans*, based on a *dpy-10* co-CRISPR protocol (Paix et al., 2015). All reagents are in 5mM Tris-HCl pH 7.5. Crispr.mit.edu and CRISPOR (http://crispor.org) web tools were used to choose a Cas9 cleavage site (NGG) close to the edit site, the best sequence of the crRNAs (50 to 75% of GC content), and for off-target prediction. 1μL of CrRNA(s) (8μg/μL or 0.6nmole/μL) and repair template(s) (1μg/μL) designed for *lgg-1* and *dpy-10* genes were mixed with 4.1 μL of *S. pyrogenes* Cas9 (20 pmole/μL) and 5 μL of universal tracrRNA (4μg/μL 4μg/μL or 0.17 nmol/μL molarity) in 0.75 μL Hepes (200 mM) 0.5μL KCl (1M) and water up to 20 μL. The mix was heated for 10 min at 37°C and injected in the gonad of young adult hermaphrodites. Progenies of injected animals were cloned and genotyped by PCR. Mutants were outcrossed and *lgg-1* gene was sequenced to check for the specific mutations.

### Nematode starvation and lifespan

For starvation experiments, adult hermaphrodites were bleached to obtain synchronized L1 larvae. L1 were incubated in 0.5 mL sterilized M9 at 20°C on spinning wheel. At each time point, an aliquot from each sample tube was placed on a plate seeded with *E. coli* OP50. The number of worms surviving to adulthood was counted 2 or 4 days after. Life span was performed on more than 100 animals for each genotype with independent duplicates and analyzes using Kaplan-Meier method and Log-Rank (Mantel-Cox) test.

### RNA mediated interference

RNAi by feeding was performed as described (Kamath et al., 2003). Fourth-larval stage (L4) animals or embryos were raised onto 1 mM isopropyl-D-β-thiogalactopyranoside (IPTG)-containing nematode growth media (NGM) plates seeded with bacteria (E. coliHT115[DE3]) carrying the empty vector L4440 (pPD129.36) as a control or the bacterial clones from the J. Ahringer library, Open Biosystem (lgg-1: JA-C32D5.9, epg-5: JA-C56C10.12).

### Western blot

The worms were collected after centrifugation in M9 and then mixed with the lysis buffer described previously (Springhorn and Hoppe, 2019) (25mM tris-HCl,pH7.6; 150mM NaCl; 1mM ethylenediaminetetraacetic acid (EDTA) 1% Triton X-100; 1% sodium deoxycholate (w/v); 1% SDS (w/v)) containing glass beads (Sigma-Aldrich 425-600um G8772100G). They were then denatured using Precellys 24 machine at 6000rpm with incubation for about 5 min twice to cool down the sample. The protein extracts are then centrifuged at 15000 rpm and separated on a NuPAGE 4%-12% Bis-Tris gel (Life Technologies, NP0321BOX). The non-specific sites are then blocked after the incubation for one hour with PBS Tween 0.1% (Tris Base NaCl, Tween20) BSA 2%. Blots were probed with, anti-LGG-1 (1:3000 rabbit Ab#3 a generous gift from T. Hoppe or 1:200 Ab#1 (Al Rawi et al., 2011), anti-LGG-2 (1:200 rabbit), anti-Tubulin (1:1000 mouse; Sigma, 078K4763) and revealed using HRP-conjugated antibodies (1: 5000 promega W401B and 1:10000 promega W4021) and the Super Signal Pico Chemiluminescent Substrate (Thermo Scientific, 34579). Signals were revealed on a Las3000 photoimager (Fuji) and quantified with Image Lab software.

### Immunofluorescence and light microscopy

Embryos were prepared for immunofluorescence staining by freeze-fracture and methanol fixation at −20°C around 30 minutes. Staining were probed with anti-LGG-1 (1;100) and anti-GABARAP (1;200). Fifty adult hermaphrodites were cut to release the early embryos on a previously poly-L-lysinated slide (0.1%). Late embryos were deposited using a flattened platinum wire and bacteria as glue. Embryos were prepared for immunofluorescence staining by freeze-fracture and methanol fixation 30 min at −20°C, incubated 40 min in 0.5% Tween, 3% BSA, PBS solution, and washed twice 30 min in 0.5% tween PBS solution. Incubation overnight at 4°C overnight with the primary antibodies anti-LGG-1(rabbit 1:100) anti-GABARAP (rabbit 1:200) (1: 200), anti-LGG-2 (rabbit 1:100) was followed by 2 washes, 2 hours incubation at room temperature with the secondary antibodies, Alexa488 and Alexa 568 (1: 1000), and two washes. Embryos were mounted in DABCO and imaged on an AxioImagerM2 microscope (Zeiss) equipped with Nomarski optics, coupled to a camera (AxioCam506mono) or a confocal Leica TCS SP8 microscope with serial z sections of 0.5 to 1 μm. Images were analyzed, quantified and processed using ImageJ or Fidji.

### Electronic microscopy

One-day adults were transferred to M9 20% BSA (Sigma-Aldrich, A7030) on 1% phosphatidylcholine (sigma-aldrich) pre-coated 200μm deep flat carriers (Leica Biosystems), followed by cryo-immobilization in the EMPACT-2 HPF apparatus (Leica Microsystems; Vienna Austria) as described (Largeau & Legouis, 2019). Cryo-substitution was performed using an Automated Freeze-substitution System (AFS2) with integrated binocular lens, and incubating chamber (Leica Microsystems; Wetzlar, Germany) with acetone. Blocks were infiltrated with 100% EPON, and embedded in fresh EPON (Agar Scientific, R1165). Ultrathin sections of 80 nm were cut on an ultramicrotome (Leica Microsystems, EM UC7) and collected on a formvar and carbon-coated copper slot grid (LFG, FCF-2010-CU-50). Sections were contrasted with 0,05% Oolong tea extract (OTE) for 30 minutes and 0.08 M lead citrate (Sigma-Aldrich, 15326) for 8 minutes. Sections were observed with a Jeol 1400 TEM at 120 kV and images acquired with a Gatan 11 Mpixels SC1000 Orius CCD camera.

### *S. cerevisiae* culture and autophagy assays

The quantitative Pho8Δ60 assay of nonspecific autophagy, was performed as described (Noda and Klionsky, 2008). Cells were grown to log phase in YPD medium then were transferred to nitrogen starvation medium for 4h. At different time point, 5 OD600 units of cells were collected, washed and resuspended in ice-cold assay buffer (250mM Tris-HCl, pH 9; 10mM MgSO4 and 10 μM ZnSO4) with 1mM PMSF. Then cells were broken using glass beads. For the assay, 10 μl of lysed cells are added to 500 μl of ice-cold assay buffer, placed at 30°C for 5 min before tadding 50 μl of 55 mM α-naphthyl phosphate disodium salt for 20 min at 30°C. The reaction was stopped with 500 μl of 2M glycine-NaOH, pH 11 and the fluorescence measured (345 nm excitation / 472 nm emission). The Pho8Δ60 activity correspond at emission per amount of protein in the reaction (mg) and reaction time (min).

The number of vacuoles was counted after incubation of exponentially growing cells with FM4-64 (33 μM) in YPD medium at 30°C for one hour, washing and imaging.

For survival to nitrogen starvation cells were grown to log phase in appropriate complete synthetic medium (CSM) and transferred to nitrogen starvation medium (0.17% yeast nitrogen base and 2% glucose). After 0 to 6 days of starvation, cells were spread on YPD plates and colonies were counted after 2 days at 30°C. For LGG-1 rescue assays, LGG-1(G116A) and LGG-1(G116AG117*) were generated by PCR amplification from cDNA LGG-1 and cloned in pRS416 vector under the control of GPD promoter.

### Affinity purification of LGG-1

1 mg of total proteins from *C. elegans* lysate were incubated on ice 10 minutes in 800μL of TUBE lysis buffer [50 mM sodium fluoride, 5 mM tetra-sodium pyrophosphate, 10 mM b-glyceropyrophosphate, 1% Igepal CA-630, 2 mM EDTA, 20 mM Na_2_HPO_4_, 20 mM NaH_2_PO_4_, and 1.2 mg/ml complete protease inhibitor cocktail (Roche, Basel, Switzerland)] supplemented with 200μg of purified LC3 traps or GST control. After cold centrifugation at 16200 g for 30 min, supernatant was harvested and added to 400μl of prewashed glutathione-agarose beads (Sigma), and incubated for 6 hours rotating at 4°C. Beads were centrifugated at 1000 g for 5 min at 4°C (Beckman Coulter Microfuge 22R, Fullerton, CA, USA), washed five times using 10 column volumes of PBS-tween 0.05%. Elution was done in 100 μL of [Tris pH7.5, 150mM NaCl, 1% Triton, 1% SDS] at 95°C during 10min, and supernatant was harvested.

### Mass spectrometry

Protein samples affinity purification were prepared using the single-pot, solid-phase-enhanced sample-preparation (SP3) approach as described (Hughes et al., 2019). Samples were mixed with 10μl of 10μg/μl solution of Sera-Mag SpeedBeadsTM hydrophilic and hydrophobic magnetics beads (GE healthcare, ref 45152105050250 and 65152105050250) with a beads to sample ratio of 10:1. After a binding step in 50% ethanol in water, and three successive washes with 80% ethanol in water, sample were digested with 100μl of a 5ng/ul sequencing grade modified trypsin solution (PROMEGA). 50μl of Trypsin-generated peptides were vacuum dried, resuspended in 10μl of loading buffer (2% acetonitrile and 0.05% Trifluoroacetic acid in water) and analyzed by nanoLC-MSMS using a nanoElute liquid chromatography system (Bruker) coupled to a timsTOF Pro mass spectrometer (Bruker). Briefly, peptides were loaded on an Aurora analytical column (ION OPTIK, 25cm x75μm, C18, 1.6μm) and eluted with a gradient of 0-35% of solvent B for 100 min. Solvent A was 0.1 % formic acid and 2% acetonitrile in water, and solvent B was 99.9% acetonitrile with 0.1% formic acid. MS and MS/MS spectra were recorded and converted into mgf files as previously described. Proteins identification were performed with Mascot search engine (Matrix science, London, UK) against a database composed of all Lgg1 sequences including the wild-type and mutant sequences. Database searches were performed using semi-trypsin cleavage specificity with five possible miscleavages. Methionine oxidation was set as variable modification. Peptide and fragment tolerances were set at 15 ppm and 0.05 Da respectively. A peptide mascot score threshold of 13 was set for peptide identification. C-terminal peptides were further validated manually.

## QUANTIFICATION AND STATISTICAL ANALYSIS

All data summarization and statistical analyses were performed by using either the GraphPad-Prism or the R software (www.r-project.org). The Shapiro-Wilk’s test was used to evaluate the normal distribution of the values and the Hartley Fmax test for similar variance analysis. Data derived from different genetic backgrounds were compared by Student t test, Anova, Kruskal-Wallis or Wilcoxon-Mann-Whitney tests. The Fisher’s exact test was used for nominal variables. Longevity was assessed using Log-Rank (Mantel-Cox) test. Boxplot representations indicate the minimum and maximum (excluding any outliers), the first (Q1/25th percentile), median (Q2/50th percentile) and the third (Q3/75th percentile) quartiles. NS (Not Significant p>0.05; * 0.05>p>0.01, **0.01>p>0.001, *** 0.001>p>0.0001 and **** p<0.0001. Exact values of n and statistical tests used can be found in the figure legends.

---

**Supplementary Table 1:**
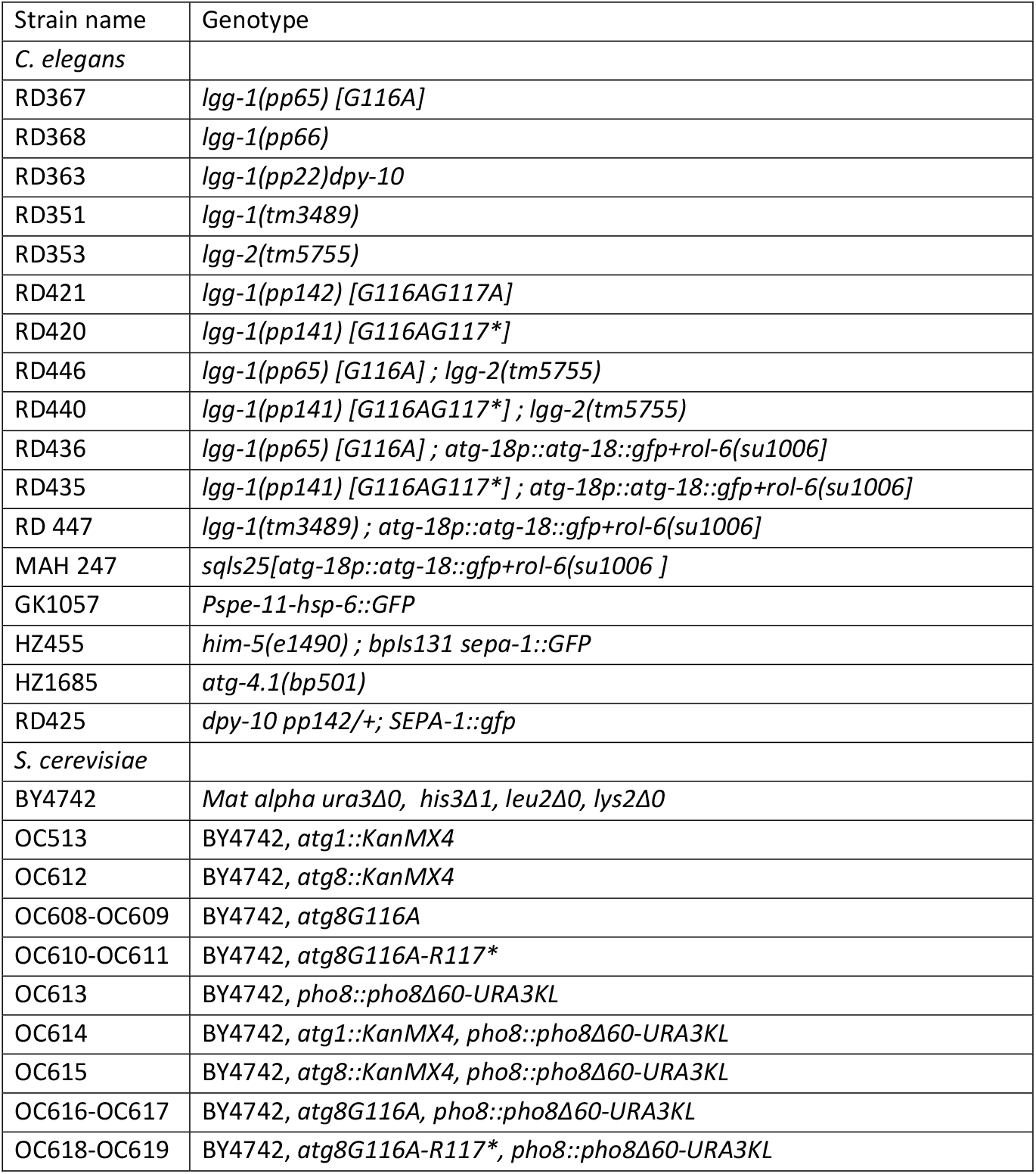
Strains used in this study.

